# Plasmodium falciparum Myosin F with a Rab-like tail domain localises to perinuclear membranes and associates with trafficking proteins

**DOI:** 10.1101/2025.11.10.687607

**Authors:** Alexander J. Holmes, Philip Ilani, Christopher Batters, Alison Kemp, Julian C. Rayner, Folma Buss

## Abstract

Members of the myosin superfamily are found in all eukaryotes, including *Plasmodium falciparum*, the parasite that causes malaria. *Plasmodium* falciparum expresses six myosins, but apart from PfMyoA, these motors remain largely uncharacterised. This includes the class XXII myosin PfMyoF. Here, we characterise PfMyoF using structural prediction tools, biochemical assays, and advanced imaging. We show that PfMyoF is a plus-end directed, processive motor with a long neck domain containing six IQ motifs that bind the calmodulin homologue PfCaM. PfMyoF can dimerise and is predicted to adopt an autoinhibited conformation via tail backfolding. The PfMyoF tail contains a Rab-like domain that has not been identified in any other known class of myosin. Pull-down experiments show interactions between endogenous PfMyoF and trafficking proteins, including the vesicle marker PfRab18. Expansion and immunoelectron microscopy reveal that PfMyoF localises to a perinuclear membrane compartment. Our findings define PfMyoF as the first known myosin with a Rab-like domain and highlight its potential role in parasite membrane trafficking pathways.

## Introduction

Myosin motor proteins are ATP-dependent mechanoenzymes which generate mechanical force through movement along actin filaments, facilitating essential cellular processes such as intracellular transport, migration and cytokinesis^1^. Given the wide range of cellular processes involving myosin motors, several myosin classes have been successfully exploited as therapeutic targets for the treatment of a variety of human disease^2^. Although the myosin superfamily is a large and diverse group of proteins, they all share a conserved, tripartite domain organisation consisting of an N-terminal motor domain that binds actin and has ATPase activity, a neck region containing between one and six IQ motifs that bind light chains or calmodulin and, with the exception of *Apicomplexan* class XIV myosins, a highly variable C-terminal cargo-binding tail domain^3^.

While the human genome contains 40 myosin genes belonging to 12 classes, *Apicomplexan* parasites such as *Plasmodium falciparum,* the causative agent of the most severe cases of malaria, possess a limited and largely uncharacterised set of unconventional myosins that are distinct in sequence and structure from those of their human host. This evolutionary divergence offers a promising opportunity to exploit *P. falciparum* myosins as selective drug targets. Malaria remains the deadliest parasitic disease worldwide. In 2025, over 40% of the world’s population are at risk of infection, with the disease estimated to kill half a million people every year. According to the most recent data, malaria was responsible for at least 597,000 deaths in 2023, 95% of which occurred in Africa^4^. While recently developed vaccines like RTS/S (Mosquirix™) have shown moderate effectiveness in reducing malaria cases (approximately a 36% reduction in young children), they do not offer complete protection, and their efficacy has been shown to decline over time^5^. In addition, resistance to existing anti-malarial drugs is an increasing problem – underscoring the urgent need for new drug targets.

The six myosins identified in *P. falciparum* include PfMyoA, PfMyoB and PfMyoE of class XIV; PfMyoF of class XXII; and PfMyoJ and PfMyoK, which belong to a class VI-like group of myosins^6^. Among these, the essential myosin motor PfMyoA has been extensively characterised and shown to function in gliding motility and red blood cell invasion, a process essential for both parasite replication and malaria pathology^7^. Furthermore, this myosin has been successfully targeted using the small molecule inhibitor KNX-002, highlighting myosin motors as suitable candidates for anti-malarial drug development^8^.

PfMyoF, a myosin of class XXII, has been shown to function during the asexual blood stage of the *P. falciparum* life cycle, the stage responsible for the clinical symptoms and mortality associated with malaria^9^. Class XXII myosins are unique to *Apicomplexan* parasites, and despite their proposed essentiality, they remain poorly characterised in terms of motor properties, interaction partners, and functional roles within the parasite.

The most detailed studies of Myosin F in *Apicomplexa* have been in *T. gondii,* where TgMyoF functions in organelle positioning and dynamics. Disruption of TgMyoF function through expression of dominant-negative constructs or conditional knockdown impairs the dynamics and distribution of organelles including dense granules (secretory organelles), the apicoplast (*Apicomplexan* metabolic plastids), and rhoptries (secretory organelles) as well as the Golgi and post-Golgi compartments positive for Rab5, Rab6, Rab7, Syn6 and DrpB^10–12^. In addition, the expression of dominant-negative TgMyoF constructs also caused centrosomal mislocalisation during cell division, indicating a role in maintaining centrosome positioning.

In *P. falciparum*, the first functional study of PfMyoF demonstrated its expression throughout the asexual blood cycle, particularly in the trophozoite and schizont stages^9^. Using a knock-sideways approach, the study showed that conditional mislocalisation of PfMyoF impaired haemoglobin uptake and parasite growth, implicating PfMyoF in endocytic processes essential for nutrient acquisition. A small fraction of PfMyoF was observed to colocalise with Kelch13-positive vesicles, structures associated with the endocytosis of host cytoplasm and potentially linking PfMyoF to artemisinin resistance; however, the C-terminal 2xFKBP-GFP-2xFKBP tag used in the study was found to affect PfMyoF function, leaving the precise localisation of endogenous PfMyoF unknown.

Genetic screens provide further evidence for the importance of class XXII myosin function in *Plasmodium* subspecies. In *Plasmodium berghei*, a rodent model of malaria, Wall and colleagues GFP-tagged all six myosins, revealing distinct patterns of expression and localisation across different life cycle stages^13^. Gene knockout experiments indicated that PbMyoA, PbMyoF and PbMyoK are essential, as deletion of these genes yielded no viable parasites. A recent high-saturation whole-genome piggyBac screen in *P. knowlesi* identified PkMyoF as an essential protein for parasite viability^14^. In *P. falciparum*, only PfMyoA has so far been identified as essential^15^. Taken together, these findings suggest that PfMyoF is likely to be essential for *P. falciparum* viability, but further targeted validation is needed.

While these studies have implicated PfMyoF in intracellular transport and parasite viability, its fundamental motor properties and cargo binding ability remain uncharacterised. To date, no studies have investigated the basic sequence and structural features of PfMyoF including its oligomerisation status, directionality along actin filaments, domain architecture, or potential binding partners and endogenous localisation. Such insights are essential to understand the mechanistic basis of PfMyoF function and in assessing its potential as a therapeutic target.

Here, we address these knowledge gaps in PfMyoF using a combination of bioinformatics, structural modelling, *in vitro* expression and cell-based assays. We identify unique sequence and domain features of PfMyoF including the discovery of a novel Rab-like domain within the tail. Using AlphaFold modelling we further highlight that the PfMyoF lever arm contains six light chain binding sites for PfCaM, and suggest that PfMyoF is likely a dimer, supporting a role in processive cargo transport. Using a novel AlphaFold-based screen and cell-based assay, we determine that PfMyoF is a plus-end directed myosin. Finally, using a polyclonal antibody against the WD40 domain, we identify putative interaction partners and the cellular localisation of the endogenous protein at a perinuclear membrane compartment. Overall, this study combines computational and experimental approaches to define the core properties of PfMyoF, laying the groundwork for dissecting its role in *P. falciparum* biology.

## Results

### PfMyoF has a Conserved Motor Domain, *Plasmodium*-Specific Insertions and a Unique Tail

To gain insight into the structural organisation of PfMyoF, we used AlphaFold to generate a full-length structural model. The predicted PfMyoF structure follows a canonical myosin organisation with the motor domain at the N-terminus, followed by a long neck domain, an extended coiled-coil region, and two unique C-terminal tail domains: a previously unannotated Rab-like domain, and finally a WD40 β-propeller (Fig 1A and B). PfMyoF adopts a compact, backfolded structure, suggestive of an autoinhibited conformation (Fig 1Ba, 1Bb and 1Bc) that resembles TgMyoF but is distinct from other classes of *Apicomplexan* myosins (Fig. S1). The model has a moderate pTM score of 0.57, indicative of a reasonably confident prediction, with higher local confidence in the motor and individual tail domains highlighted by darker green in the Predicted Aligned Error (PAE) plot (Fig 1Bb).

**Fig 1.**
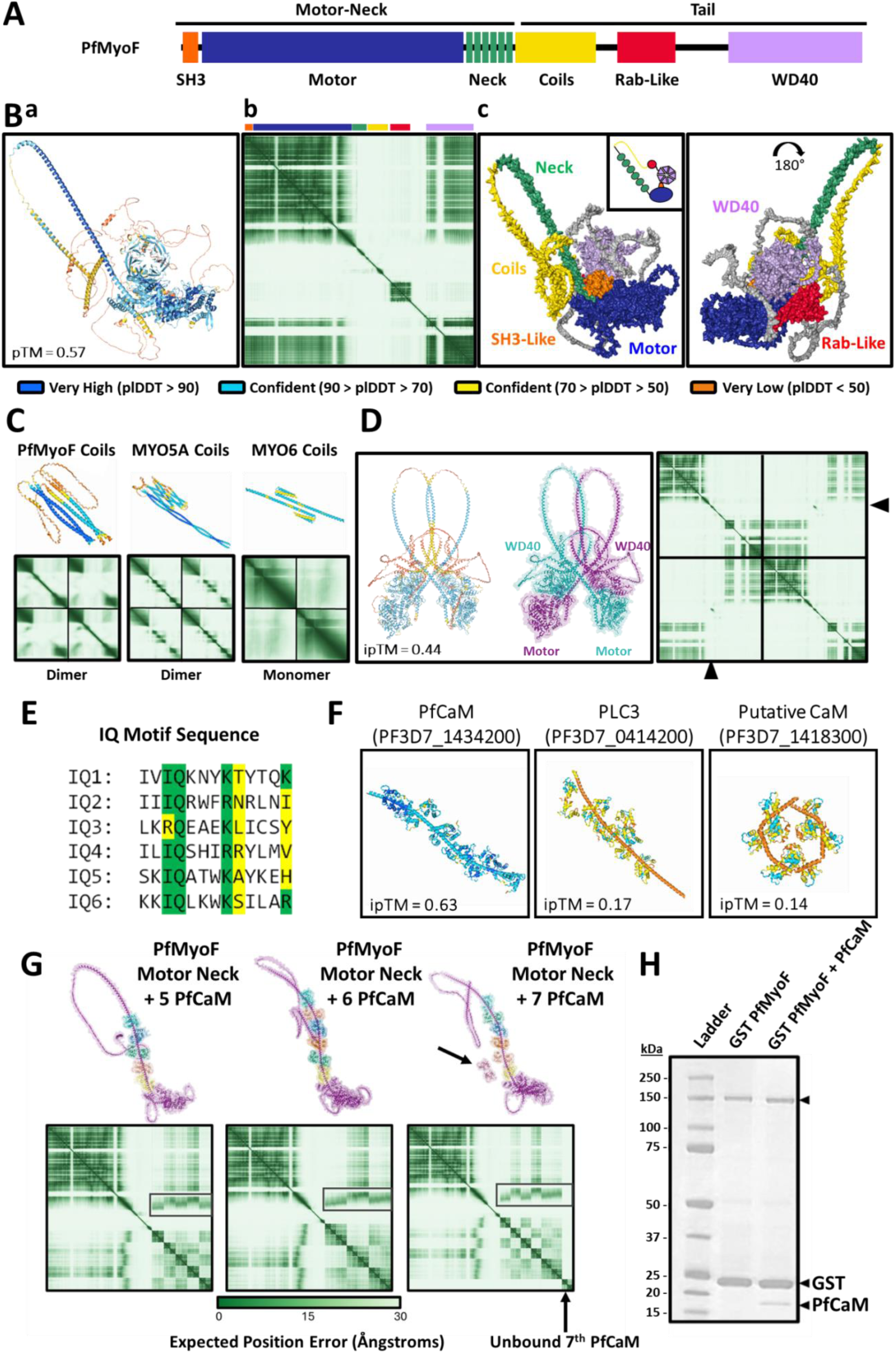
Predicted structural features and dimerisation potential of PfMyoF. (A) Schematic of the full-length PfMyoF, highlighting the N-terminal motor domain, neck with six IQ motifs, coiled-coil regions, Rab-like domain, and C-terminal WD40 domain. (B) AlphaFold-3-predicted structure based on the PfMyoF sequence. (Ba) PfMyoF coloured by pLDDT value. (Bb) PAE plot of PfMyoF. (Bc) PfMyoF coloured according to key domains. (C) AlphaFold-3-predicted structures of the coil domains of PfMyoF, HsMYO5A (positive control) and HsMYO6 (negative control). Structures are coloured by pLDDT with corresponding PAE plots presented below. (D) AlphaFold-3 prediction of PfMyoF dimers show coiled-coil interaction with a backfolded architecture. Black arrowheads point to the coiled-coil interaction within the PAE plot. (E) Sequence alignment showing the six putative IQ motifs within PfMyoF neck domain. Green sequences highlight the canonical IQ motif amino acids and yellow sequence show degenerate sequences. (F) AlphaFold-3 prediction of putative light chains PfCaM, PLC3 and PF3D7_1418300 show that only PfCaM binds to all six IQ motifs in the PfMyoF neck with high ipTM confidence (0.63). (G) AlphaFold-3 modelling predicts binding of up to six PfCaM molecules along the PfMyoF neck domain (purple) ending at the beginning of the coil domain. Additional PfCaM proteins remain unbound (indicated on the PAE plot by a black arrow). (H) SDS-PAGE gel highlighting co-purification of 6His-PfCaM with GST-PfMyoF motor neck from Sf9 cells, confirming its interaction with PfMyoF.

The motor domain of PfMyoF shares core architectural features with other myosin classes, including the essential conserved ATP-binding P-loop (GESGAGKT), the actin-binding interface, and a preserved cleft architecture (Fig 1B & Fig. S2A). Other key myosin features including the TEDS site, which is a regulatory phosphorylation site common to unconventional myosins, is also conserved. PfMyoF, like all *P. falciparum* myosins, contains an N-terminal SH3-like domain (Fig S2A). In human myosins MYO5A and MYO6, this domain interacts with the tail to stabilise the folded autoinhibited conformation^16,17^. Unlike the well-characterised PfMyoA, which contains a phospho-regulated S19 residue preceding its SH3-like domain, PfMyoF lacks this extension and phosphorylation motif^18^.

The *Plasmodium* genome is highly AT-rich, and the PfMyoF sequence reflects this bias, with an overall AT content of ∼75% (in contrast to human myosins like MYO5A with an AT content of 55%). This compositional skew likely contributes to the presence of large, asparagine-rich insertions not observed in myosins from other organisms such as *Toxoplasma* TgMyoF (Fig 1Ba and 1Bc, S2A, S2B and S2C). These insertions are predicted to be unstructured and may influence folding, solubility, or regulation.

Together, these findings indicate that PfMyoF retains a structurally conserved core motor domain but has evolved *Plasmodium*-specific insertions and unique tail domains, which may be central to its parasite-specific functions and represent potential therapeutic targets.

### AlphaFold-3 prediction indicates that PfMyoF has the potential to form dimers

Myosins can be broadly classified based on their oligomeric state into monomeric (single-headed) or dimeric (double-headed) motors such as myosins of class V. Dimerisation, which enables hand-over-hand, processive movement along actin filaments, typically occurs via coiled-coil formation in alpha helical regions within their tail domains. The coiled-coil prediction software MARCOIL identifies two coiled-coil regions in the PfMyoF tail^19^. To assess their dimerisation potential, we used AlphaFold-3, which accurately distinguishes between the dimeric MYO5A and the monomeric MYO6: MYO5A forms as a coiled-coil dimer whereas MYO6 contains a single alpha helix (SAH domain) that does not dimerise (Fig 1C)^20^. Similarly to MYO5A, AlphaFold-3 predicted that two copies of PfMyoF can form a dimer mediated by coiled-coil interactions (Fig 1C). The contribution of these coiled-coil regions to PfMyoF or MYO5A dimerisation is highlighted by the PAE plots (Fig 1C). Furthermore, the full-length dimeric PfMyoF is predicted to adopt an autoinhibited conformation with the tail domains back-folding to interact with the motor domains, as observed for MYO5A (Fig 1D)^21^.

### PfMyoF contains six light chain binding sites and a capacity to bind PfCaM

Next, we focused on identifying the PfMyoF light chains, which are critical for motor activation. The neck domain, together with the converter region, forms the swinging lever arm that drives myosin movement upon nucleotide exchange^22^. Calmodulin-like light chains bind to IQ motifs within the neck, stabilising and regulating the swinging lever arm motion. To date, only the light chains of PfMyoA and PfMyoB from *P. falciparum* have been identified. PfMyoA binds the MTIP (PF3D7_1246400) and ELC (PF3D7_1017500) light chains^23,24^, whereas PfMyoB binds to MLC-B (PF3D7_1118700) and potentially other light chains^25^. While many human unconventional myosins complex with calmodulin, neither class XIV myosin appears to bind the *P. falciparum* calmodulin homologue (PfCaM).

Sequence analysis of PfMyoF identified six degenerate IQ motifs broadly matching the consensus sequence ‘IQxxxRQxxxR’, suggesting the presence of six putative light chain binding sites (Fig 1E). In addition to PfCaM, which shows 89% sequence conservation with human CaM, nine other calmodulin-like proteins were identified based on sequence identity, and all were tested for their ability to bind to the PfMyoF neck domain using AlphaFold-3 (Fig 1F). Modelling with six PfCaM copies yielded an ipTM score of 0.63, which exceeds the 0.6 threshold for confident interaction, supporting a plausible linear lever arm structure (Fig 1F, left panel). In contrast, all other CaM-like proteins gave lower ipTM scores (ranging from 0.13 to 0.36) indicating weaker or unlikely interactions (Fig 1F, middle and right panel).

Myosins with six IQ motifs, such as human MYO5A, typically bind multiple copies of a single light chain. AlphaFold-3 modelling of PfCaM binding to the neck region of PfMyoF showed that PfCaM binds sequentially to all six IQ motifs, saturating at six molecules with no additional binding (Fig 1G).

The AlphaFold-3 prediction was verified experimentally by co-expressing GFP-tagged PfMyoF motor-neck alongside 6His-PfCaM in Sf9 cells (Fig 1H). Small quantities of soluble GFP-PfMyoF motor-neck could be purified using GST-GFP nanobody bound to glutathione Sepharose and analysed by SDS-PAGE. In Sf9 cells expressing only PfMyoF, no band is present at the expected kDa of a light chain such as PfCaM. However, in cells co-expressing PfMyoF and PfCaM, an additional 18 kDa band is present at the predicted size for 6His-PfCaM (Fig 1H). This co-purification of PfCaM with PfMyoF provides direct evidence that PfCaM can function as a PfMyoF light chain.

### The unique Rab-like Domain of PfMyoF

The functional differences between classes of unconventional myosins are determined by their unique tail domains, which contain distinct motifs and binding sites. As such, the tail domain determines both the cellular localisation and binding partners of the myosin, allowing for anchoring, scaffolding, or transportation functions.

The PfMyoF tail domain contains at its C-terminus a WD40 domain, as previously described^6^. Structural predictions revealed a second previously undefined domain between the coiled-coil and WD40 domains, which shows a strong similarity to Rab proteins of the Ras family (Fig 2A and Fig S3). Rab proteins are activated by GTP binding, enabling the recruitment of trafficking machinery to vesicles and organellar membranes^26^. A Rab-like domain has not been identified in any myosin tail to date, raising questions about its function and potential for GTPase activity within the PfMyoF tail.

**Fig 2.**
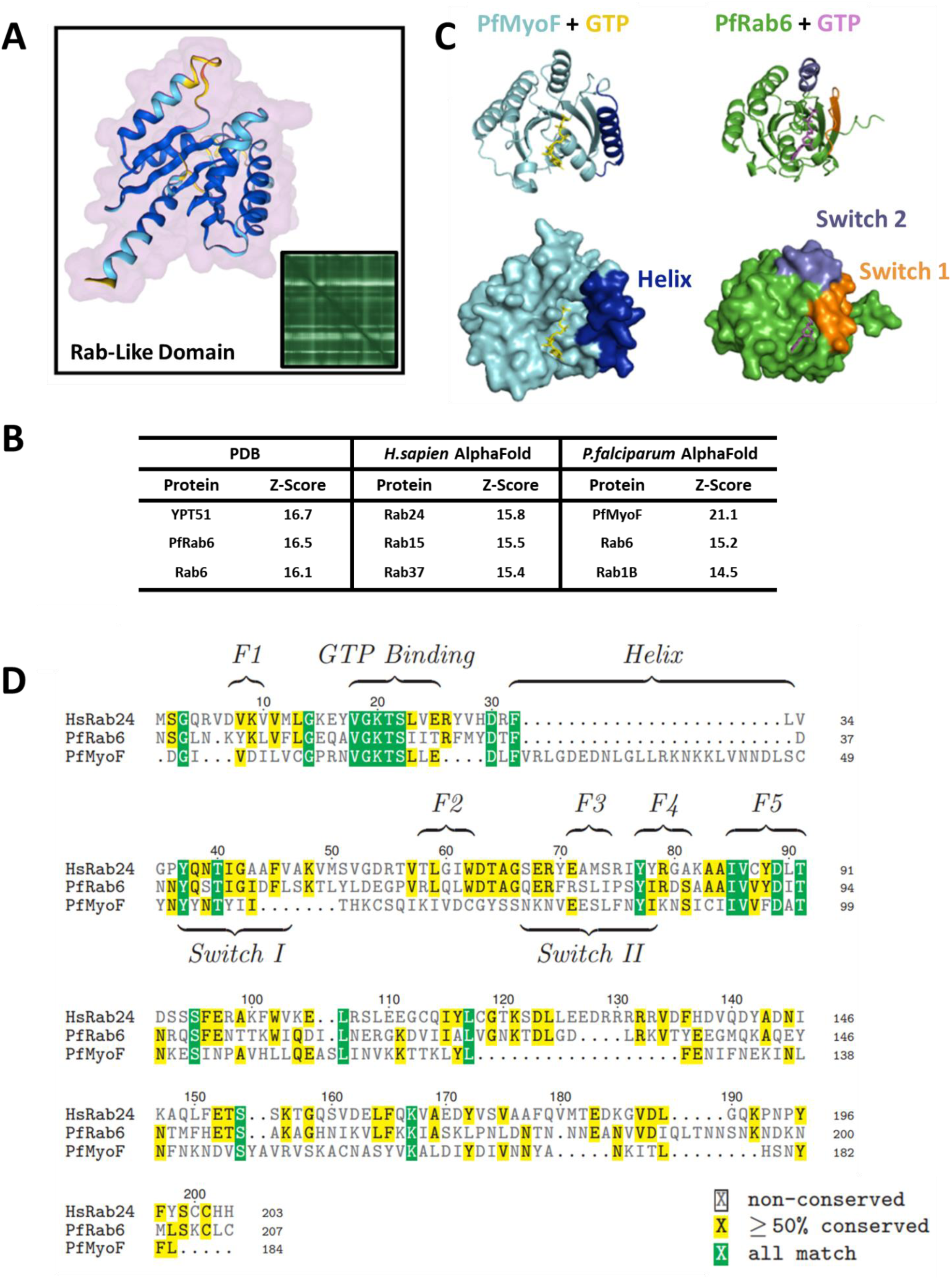
Identification and structural characterisation of the PfMyoF Rab-like domain. (A) AlphaFold-3 predicted structure of PfMyoF Rab-Like domain. (B) DALI structural homology search identifies significant similarity to *Plasmodium* and human Rab GTPases. A table detailing the top hits from a DALI server search of the protein data bank (PDB) and AlphaFold servers for proteins with a similar structure of PfMyoF Rab-Like domain. Z-scores, a measure of significant similarities, are shown. (C) AlphaFold-3 predicted structure of PfMyoF Rab-Like domain and PfRab6 bound to GTP. The additional helix in PfMyoF appears to disrupt the GTP binding pocket and reposition the key Switch I and Switch II sites essential for GTP hydrolysis. (D) Sequence alignment of PfMyoF Rab-Like domain, PfRab6 and HsRab24 reveals a conserved GTP-binding site but divergence in Switch I and II regions. Residues are coloured by conservation, and regions of interest are labelled.

Sequence homology searches for this PfMyoF Rab-like domain using BLAST^27^ identified the 11 *P. falciparum* Rab proteins as the top hits, with the Golgi-localised PfRab6 ranking highest. Structural homology modelling using the DALI server further compared the AlphaFold-predicted structure of this domain to known protein structures in the protein data bank (PDB) and other AlphaFold-predicted protein structures^28^. All the top hits (ordered by Z-score, a measure of significant similarity) from both comparisons indicated strong homology to Rab family proteins (Fig 2B).

Overlay of the AlphaFold-3-predicted structures of PfRab6 and PfMyoF Rab-like domains reveals high structural similarity. PfMyoF possesses an additional α-helix that is conserved across *Apicomplexan* Myosin F proteins (Fig 2C, helix highlighted in dark blue). Further sequence alignment with human Rab24 and PfRab6 shows conservation of the GTP-binding site in PfMyoF, but poor conservation of the Switch I and Switch II regions required for γ-phosphate binding and hydrolysis, suggesting that Rab-like domain of PfMyoFis a pseudodomain that may not possess intrinsic GTPase activity (Fig 2D). Structural alignment indicates that the additional α-helix results in the rearrangement and opening of the GTP binding cleft, and displacement of the switch regions away from the γ-phosphate of GTP (Fig 2C).

### The WD40 Domain of PfMyoF

The second of the two unique tail domains is a WD40 domain (Fig 3A), which typically assembles multi-protein scaffolds. WD40 domains consist of repeating WDR motifs that form the ‘blades’ of a rounded β-propeller. Sequence analysis and AlphaFold prediction indicate that the PfMyoF WD40 domain contains 7 blades, including a large 58-residue unstructured insertion between blades 1 and 2, specific to *Plasmodium* species (Fig 3B).

**Fig 3.**
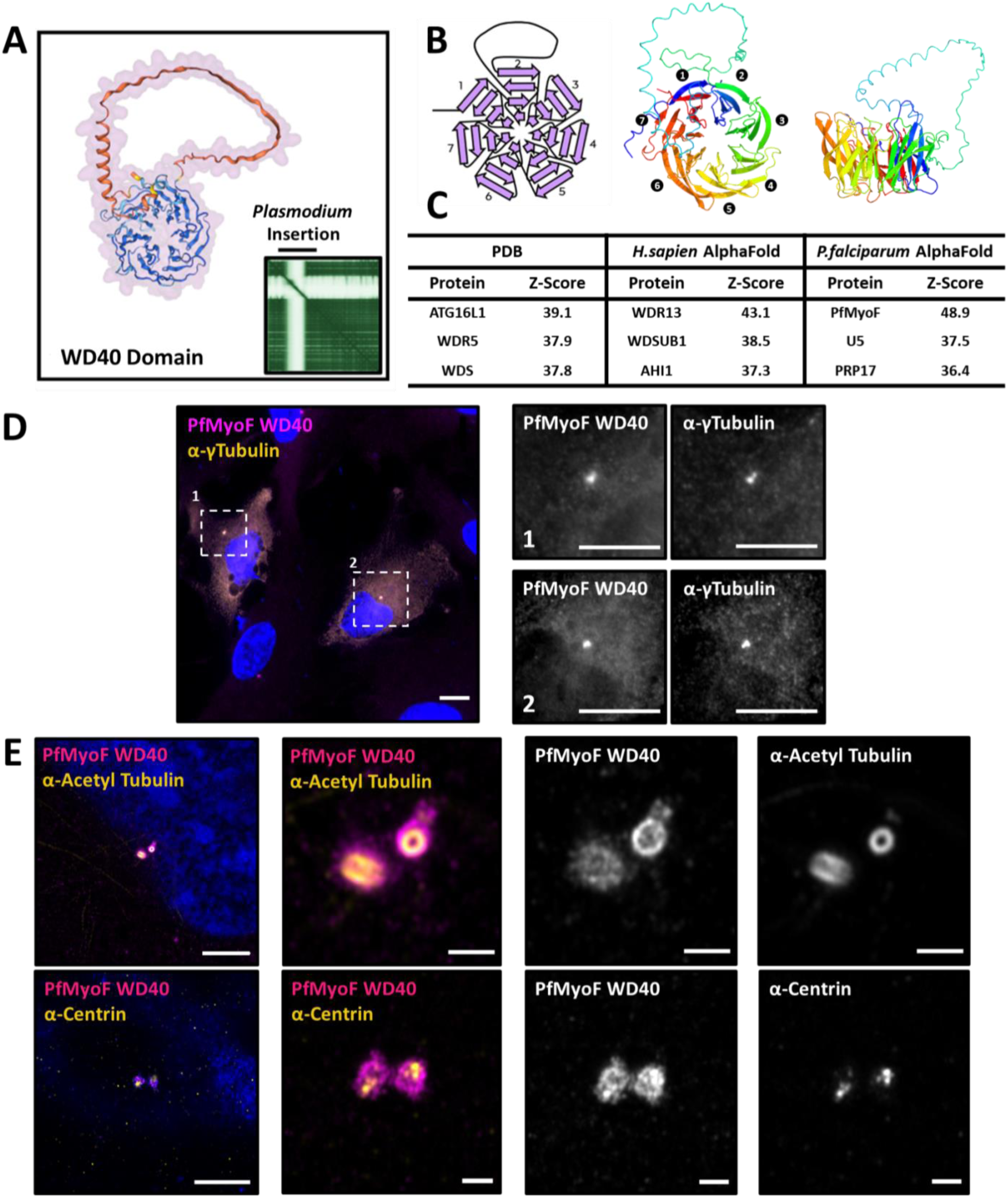
Characterisation and centrosomal localisation of the PfMyoF WD40 domain in mammalian cells. (A) AlphaFold-3 predicted structure of PfMyoF WD40 domain, forming a 7-bladed β-propeller structure with a *Plasmodium* specific insertion. (B) A schematic of PfMyoF WD40 (left) highlighting the 7-blade structure as well as an N-terminal to C-terminal rainbow colour scheme in top view (middle) and side view (right). (C) A table detailing the top hits from a DALI server search of the protein data bank (PDB) and AlphaFold servers for proteins with a similar structure of PfMyoF WD40 domain. Z-scores, a measure of significant similarities, are shown. (D) Confocal microscopy of GFP-tagged PfMyoF WD40 expressed in RPE cells stained with anti-GFP (magenta), anti-γTubulin (yellow) and Hoechst (blue). Scale bars represent 10 µm. (E) iU-ExM of RPE cells transiently expressing GFP-tagged PfMyoF WD40 domain and stained with anti-GFP (magenta), Hoechst (blue) and either anti-Centrin or anti-AcetylTubulin (yellow). Scale bars represent 10 µm, or 2 µm for zooms.

To further classify and characterise the PfMyoF WD40 domain, sequence and structural homology searches were performed using BLAST and DALI servers (Fig 3C). DALI analysis identified similar structures within diverse proteins, including human WDR5 (histone-related), WDR13 (transcription related), and AHI1 (Jouberin), which interacts with GTPases such as Rab8A to mediate the recruitment of Rabs and membrane trafficking machinery to the basal body of primary cilia. In the *P. falciparum* AlphaFold database, top hits included PRP17, among other spliceosome components. This functional diversity between homologous proteins warrants further investigation into the evolutionary and functional origins of the WD40 domain. While WD40 domains have not been observed in higher eukaryotic myosin tail domains, *Apicomplexan* parasites have evolved multiple myosin classes that contain them, including PfMyoF and PfMyoJ in *P. falciparum*, as well as TgMyoF, TgMyoH, TgMyoJ, and TgMyoL in *T. gondii* (Fig S1). These WD40 domains show considerable sequence and structural divergence, suggesting independent adaptation and distinct functions across classes.

Previous studies in *T. gondii* found that TgMyoF associates transiently with the centrosome during cell division. As many WD40 domain-containing proteins have been shown to localise to the centrosome in several species, the GFP-tagged PfMyoF WD40 domain was transiently transfected into human retinal pigment epithelial (RPE) cells and its localisation visualised. Our results show that GFP-tagged PfMyoF WD40 strongly colocalised with γ-tubulin at a single punctum resembling the centrosome (Fig 3D). Iterative Ultrastructural Expansion microscopy (iU-ExM) showed that the PfMyoF WD40 domain localises to the area surrounding acetylated-tubulin and centrin, which form a core component of the centrosome (Fig 3E)^29^. This observation suggests potential binding of the WD40 domain to the outer microtubules or components of the pericentrosomal material.

### PfMyoF is a plus-end directed myosin motor

Myosin motors can either walk towards the plus or the minus end of actin filaments, and this directionality is crucial for their function in organelle positioning, cargo transport, and actin organisation. For example, reversing the direction of class VI myosin (MYO6) motility so that it adopts a plus-end directed (MYO6+) phenotype results in a dramatic relocalisation of APPL1-positive signalling endosomes^30^. Although *P. falciparum* myosins PfMyoJ and PfMyoK have been categorised as ‘class VI-like’, suggesting they may be minus-end directed actin motors, the directionality of PfMyoF remains unknown^6^.

A crucial determinant of minus-end directed motility is Insert 2, located at the C-terminus of the converter region within MYO6, which reorients the lever arm swing^31^. Multiple sequence alignments reveal that PfMyoF lacks any Insert 2-like sequence, whereas both PfMyoJ and PfMyoK contain an insertion at the C-terminal end of their converter domain (Fig S4). The PfMyoJ insert shows strong homology with Insert 2 of MYO6 (28% identical and 47% similarity), higher than that of PfMyoK (12% identical and 35% similarity) (Fig S4).

Although PfMyoF lacks Insert 2 adjacent to the converter domain, it contains a unique large insertion *within* the converter domain (Fig 4A). To investigate whether this PfMyoF converter insertion potentially alters motor directionality along actin filaments, we developed an AlphaFold-3-based approach to examine the impact of sequence insertions on lever-arm angle. Alignment of the converter-neck regions from all human unconventional myosins, correctly distinguished plus-ended from minus-ended motors (Fig S4). Applying this method to PfMyoA, PfMyoF, PfMyoJ, and PfMyoK along with TgMyoF (which lacks the PfMyoF converter insert), revealed that PfMyoA, PfMyoF and TgMyoF converter-necks align with plus-ended myosins, whereas PfMyoJ and PfMyoK align with minus-ended motors (Fig 4B). A MYO6+ chimera lacking Insert 2, aligned with plus-end motors, consistent with experimental data^30^ (Fig 4B). Together, these sequence and structural predictions suggest that PfMyoF is a plus-end directed motor, while PfMyoJ and PfMyoK exhibit characteristics of minus-ended myosins.

**Fig 4.**
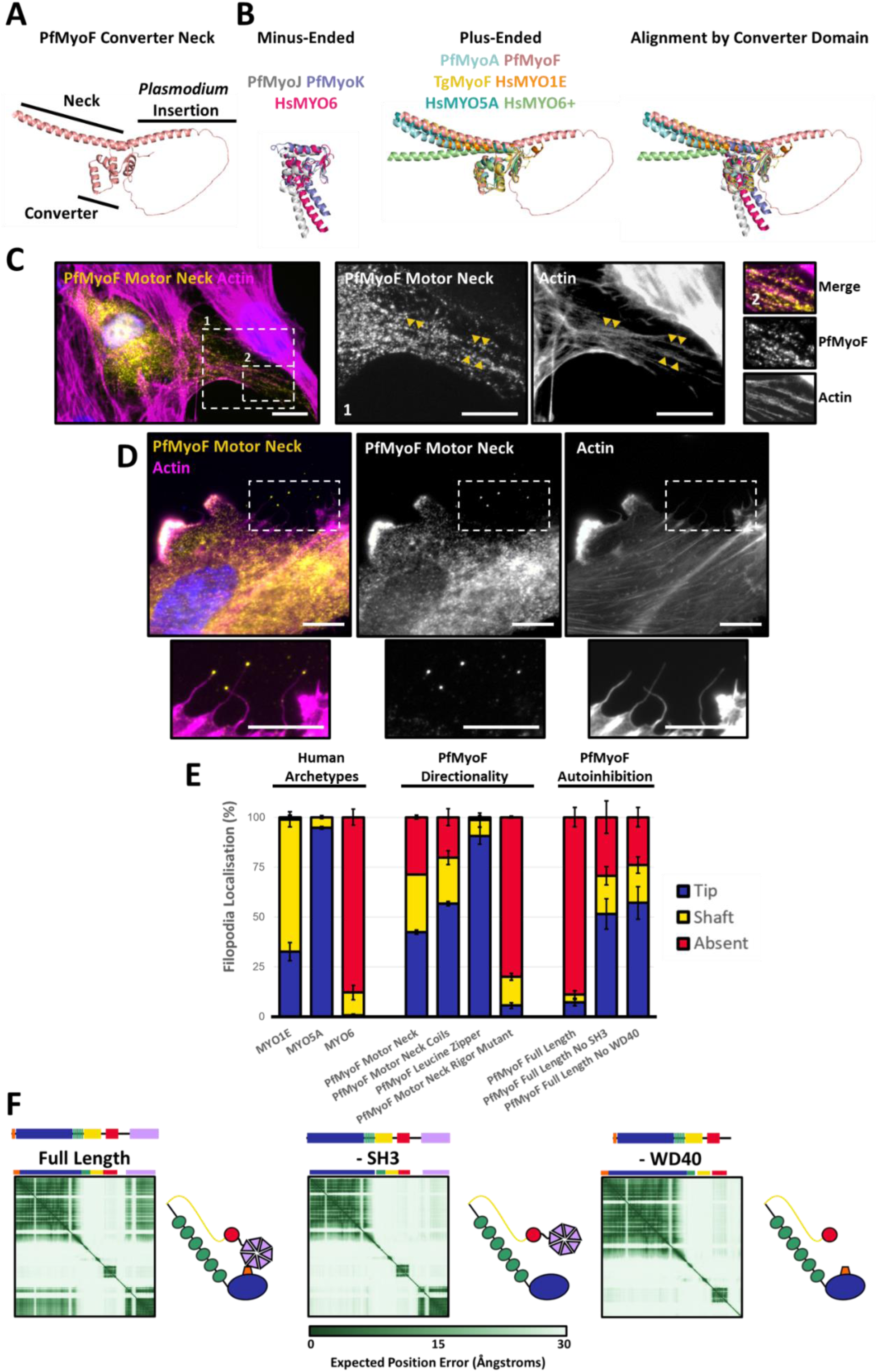
PfMyoF is a plus-end directed myosin motor. (A) AlphaFold-3 prediction of the PfMyoF converter-neck region highlights a larger insertion within the converter domain. (B) Alignment of HsMYO6 with PfMyoJ and K (left panel) and alignment of PfMyoA, PfMyoF, TgMyoF, HsMYO1E, HsMYO5A, HsMYO6+ by their converter domain (middle panel). In the right panel the overlay of both minus-ended (left panel) and plus-ended subgroups (middle panel) are shown. (C) PfMyoF aligns along actin filament bundles and accumulates in membrane ruffles in RPE cells transiently transfected with GFP-PfMyoF motor neck; GFP (yellow), Phalloidin (magenta) and Hoechst (blue). Scale bars represent 10 µm. (D) PfMyoF localises to filopodia tips. RPE cells were transfected with GFP-tagged PfMyoF motor neck and mCherry-tagged MYO6+ and stained with anti-GFP (yellow), Phalloidin (magenta) and Hoechst (blue). Scale bars represent 10 µm. (E) Quantification of filopodia tip localisation as a measure of their directionality across different human myosin classes and PfMyoF constructs. RPE cells were co-transfected with various GFP-tagged myosins alongside mCherry-tagged MYO6+. Per construct, 100 filopodia were counted in 3 independent experiments (n=3) and classified as either ‘tip’, ‘shaft’ or ‘absent’ localisation. These experiments were performed blinded. Error bars indicate SEM. (F) AlphaFold-3 predicts autoinhibited backfolding of PfMyoF via interaction between SH3-like and WD40 domain. Shown are the PAE plots and corresponding protein models of PfMyoF full-length, delta SH3-like or delta-WD40, which suggest that autoinhibition may be mediated through the SH3-like and WD40 domains.

To experimentally validate these AlphaFold predictions in a biological system, we expressed a GFP-tagged PfMyoF motor-neck construct in RPE cells, where it localised along actin filaments and accumulated in actin-rich membrane ruffles, indicating correct folding and actin binding in mammalian cells (Fig 4C). To set up the directionality assay in RPE cells, we expressed a MYO6+ chimera to induce long filopodia, in which the plus end of actin filaments point toward the filopodia tip. Different myosin classes localise to distinct regions within filopodia according to their intrinsic motor properties (Fig S5). Dimeric and processive plus-ended myosins, such as MYO5A, accumulate at the tips of the filopodia, while monomeric plus-ended myosins with membrane-binding capacity, such as those of class I, are evenly distributed along the filopodial shaft. In contrast, minus-ended myosins like MYO6 are excluded from filopodia and accumulate at the base. This assay, validated with *C. elegans* myosins of known directionality, is suitable for testing myosins from lower eukaryotes^32^.

In RPE cells, GFP-tagged PfMyoF motor-neck localises to filopodia tips, a hallmark of plus-end directed, processive motors (Fig 4D). Inclusion of the PfMyoF coiled-coil region increased tip localisation from 42% to 56% likely due to dimer formation via the coiled-coil domain which enhances processivity (Fig 4E). A C-terminal leucine zipper, which induces dimer formation, further increased PfMyoF tip localisation to 90%, confirming the importance of dimerisation (Fig 4E). In contrast an ATPase-deficient ‘rigor’ mutation (K_415_R) previously shown to inhibit MYO6 activity, abolished PfMyoF tip localisation (5%) (Fig 4E).

In contrast to the GFP-PfMyoF motor-neck construct, the full-length PfMyoF showed minimal tip localisation (7%) possibly due to misfolding, targeting of the tail domain to other cellular sites, or autoinhibition (Fig 4E). AlphaFold-3 predicts both monomeric and dimeric full-length PfMyoF to adopt a backfolded, autoinhibited conformation mediated by interactions between the N-terminal SH3-like domain and the WD40 domain (Fig 4F). Deleting either the SH3-like or WD40 domains restored PfMyoF tip localisation to 51% and 57%, respectively, matching the PfMyoF motor-neck-coiled-coil construct (Fig 4E).

Overall, these results validate a cell-based assay for testing myosin directionality, confirming PfMyoF as a plus-end directed motor, and demonstrating its capacity for dimerisation and autoinhibition.

### Characterisation of PfMyoF interactome via co-immunoprecipitation reveals link to membrane trafficking proteins

To investigate PfMyoF expression, localisation and to identify potential interaction partners, we generated a polyclonal antibody against a 20 kDa fragment of the PfMyoF WD40 domain. The PfMyoF antibody, α-PfMyoF, was validated by immunoblotting against recombinant 6His-tagged WD40 antigen expressed in Sf9 cells and by detection of endogenous PfMyoF in *P. falciparum* blood-stage lysates (Fig 5A). Lower molecular weight bands likely reflect PfMyoF degradation products arising during sample processing.

**Fig 5.**
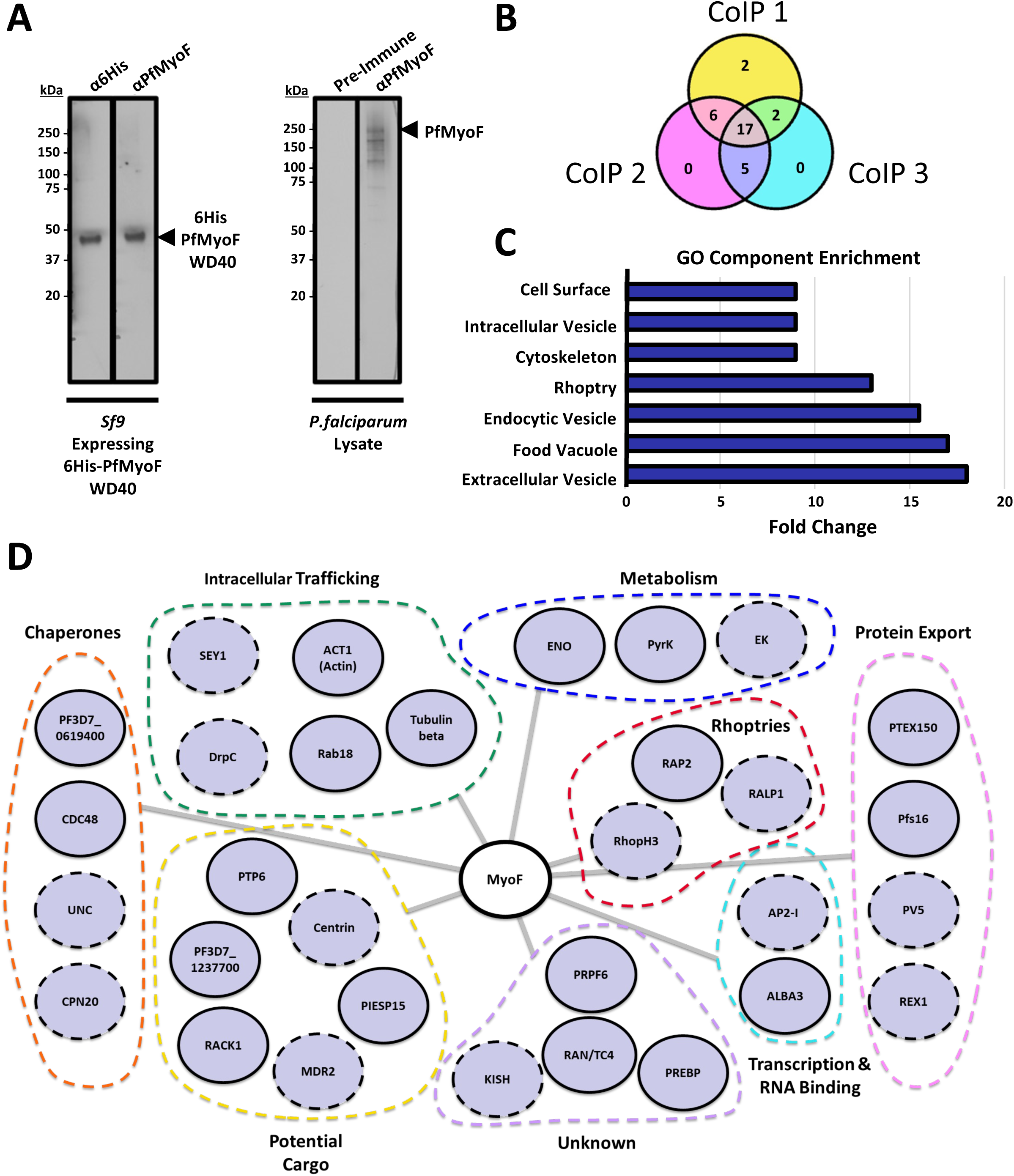
PfMyoF interactome identified by native co-immunoprecipitations. (A) Immunoblot validation of α-PfMyoF antibody against recombinant WD40 fragment and endogenous PfMyoF from parasite lysates (left panels). Sf9 cell lysate expressing 6His-WD40 domain was probed with anti-6His (left lane) or anti-PfMyoF antibodies (right lane). (Right panel) *P. falciparum* lysate was probed with Pre-Immune or anti-PfMyoF antibodies. (B) Co-immunoprecipitation from trophozoite-stage lysates identifies 32 enriched proteins; 17 were reproducible across three replicates. (C) Gene Ontology (GO) analysis of the Co-IP hits shows enrichment in vesicle-mediated transport, food vacuole, cell surface, cytoskeleton and rhoptry-associated proteins. (D) A map of proteins enriched in the anti-PfMyoF Co-IP. Hits were defined as a 5-fold enrichment in iBAQ scores of anti-PfMyoF relative to the Pre-Immune control. Hits are arranged based on their predicted functions within the parasite.

We next performed co-immunoprecipitations (Co-IP) from late trophozoite lysates using α-PfMyoF and preimmune serum as a control. PfMyoF was consistently precipitated in three independent experiments, confirming antibody specificity. Proteins enriched ≥5-fold, using intensity-based absolute quantification (iBAQ) scores as a measure of abundance relative to controls, were considered candidate interactors. This yielded 32 proteins, 17 of which were reproducibly detected across all three replicates (Fig 5B). Gene Ontology analysis revealed enrichment of proteins involved in cytoskeletal organisation and membrane trafficking, including components of the endomembrane system such as the food vacuole and intracellular vesicles (Fig 5C).

A map of all the proteins found in the PfMyoF Co-IP is shown in Fig 5D. The strongest hit, PfRab18, is involved in early secretory trafficking and associates with vesicles at the ER, Golgi, and the parasitophorous vacuole membrane^33^. Two dynamin superfamily proteins, DrpC, a mitochondrial fission factor, and SEY1, an ER tubule-forming GTPase, were also identified. Furthermore, the cytoplasmic AAA+ ATPase CDC48 and its paralog PF3D7_0619400 at the apicoplast membrane, were both enriched, suggesting a potential role for PfMyoF in ER-associated degradation (ERAD) and apicoplast-specific trafficking. Although less consistently detected, other candidates include centrin, a centrosome marker.

### Endogenous PfMyoF closely associates with Golgi membranes in the perinuclear area

To further investigate the potential role of PfMyoF in intracellular transport, the α-PfMyoF antibody was used to determine the endogenous localisation of this myosin during *P. falciparum* blood-stage development. Given the small size of *P. falciparum* parasites, Ultrastructural Expansion Microscopy (U-ExM) was performed to enhance resolution^34^. In trophozoite stage parasites, the antibody to PfMyoF revealed a predominantly perinuclear staining, enriched in a discrete region that partially colocalises with the endoplasmic reticulum marker BiP (Fig 6A). In addition, a weak speckle-like distribution throughout the cytoplasm was observed. A similar staining pattern was also present in schizont-stage parasites, with one PfMyoF-enriched perinuclear dot per schizont and a speckle-like distribution in the cytoplasm (Fig 6B). Among the limited set of *P. falciparum*-specific antibodies available, the PfMyoF perinuclear site staining most closely resembled that of the cis-Golgi marker ERD2 (Fig 6C). Staining with preimmune serum yielded no specific localisation within parasites, with only low-level background signal at the parasite surface and the surrounding red blood cells (Fig 6D).

**Fig 6.**
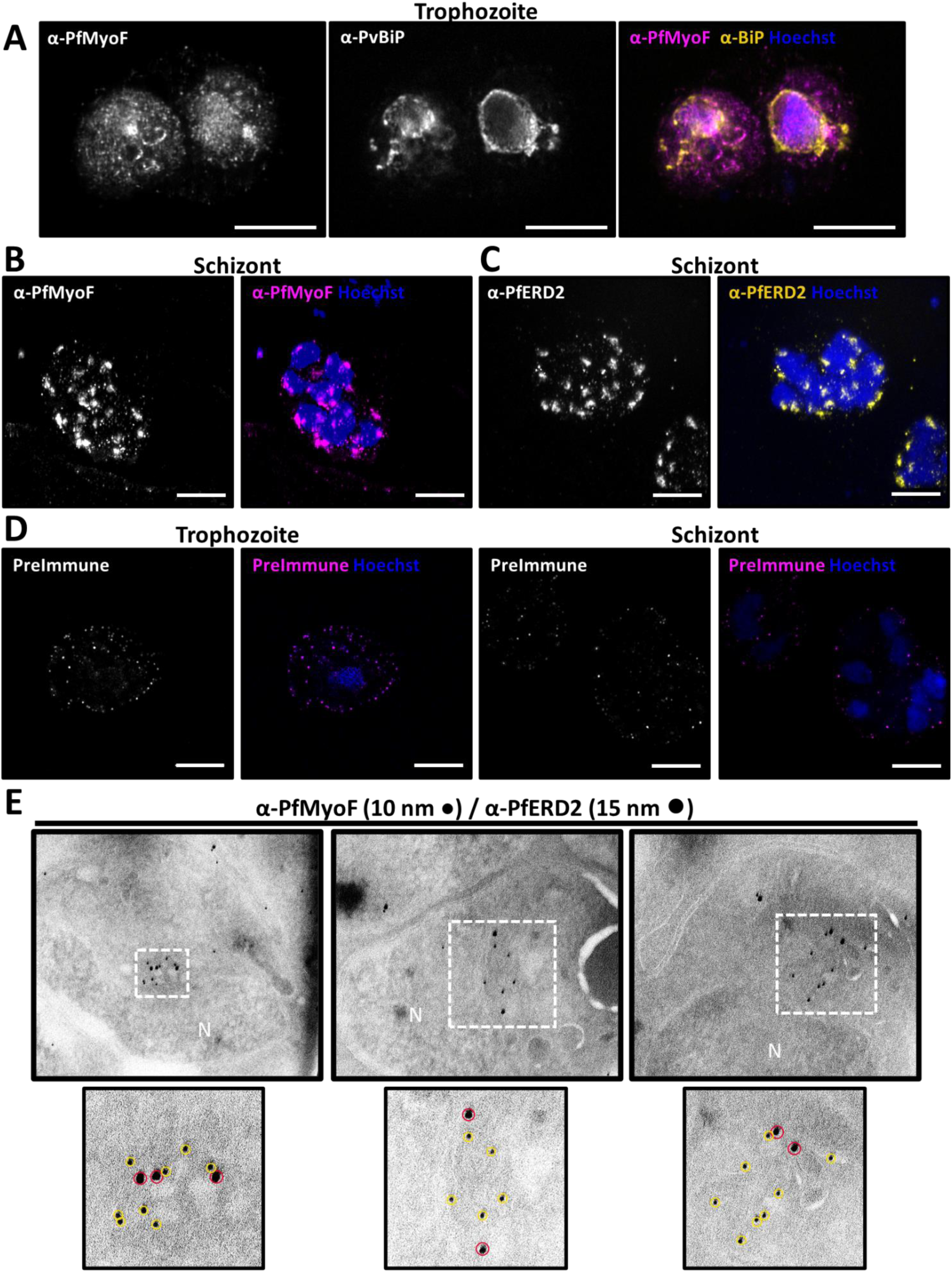
PfMyoF localises to perinuclear region around the Golgi apparatus in *P. falciparum* parasites. – (A) U-ExM of trophozoite stage parasites stained with anti-PfMyoF (magenta), the ER marker anti-PvBiP (yellow) and Hoechst (blue) shows enriched of PfMyoF near the nucleus partially overlapping with BiP (ER marker). (B) In U-ExM of schizont stage parasites stained with anti-PfMyoF (magenta) and Hoechst (blue) PfMyoF remains concentrated in the perinuclear region. (C) U-ExM of schizont stage parasites stained with anti-PfERD2 (magenta) and Hoechst (blue) shows that PfMyoF localisation closely resembles the cis-Golgi marker ERD2. (D) U-ExM of a trophozoite (left) and schizont stage parasites (right) stained with PfMyoF Pre-Immune serum gives no specific staining. (E) Immunogold labelling of schizont stage parasites with anti-PfMyoF detected with Protein A conjugated 15nm gold and anti-PfERD2 detected with Protein A conjugated 10 nm gold beads confirms co-localisation of PFMyoF and ERD2 on Golgi-proximal membranous structures. Red and yellow circles highlight 15 nm and 10 nm gold particles respectively. All scale bars represent 10 µm.

As the α-PfMyoF and α-ERD2 antibodies were both raised in rabbits, colocalisation studies using conventional fluorescence or expansion microscopy were not feasible. We therefore performed immunogold electron microscopy to assess the colocalisation between PfMyoF and ERD2. Schizont stage parasites were embedded in gelatin, snap-frozen, and cut into ultrathin sections. These were sequentially labelled with α-PfMyoF detected with Protein A - 15 nm gold conjugates followed by α-ERD2 detected with Protein A - 10 nm gold conjugates. In many parasites, PfMyoF (15 nm gold) and ERD2 (10 nm gold) were observed in close proximity on membranous structures adjacent to the nucleus (Fig 6E). As ERD2 is a marker of the parasite Golgi complex, these results suggest that PfMyoF is present in the perinuclear region in close vicinity to the Golgi apparatus in *P. falciparum.* The additional cytoplasmic PfMyoF signal may correspond to Golgi-derived or Golgi-destined vesicles, although this remains to be confirmed.

## Discussion

This study provides the first comprehensive characterisation of PfMyoF, a divergent class XXII myosin in *P. falciparum*, identifying structural as well as functional features. We demonstrate that PfMyoF is a plus-end directed, processive motor with a potential to dimerise and to adopt a folded autoinhibited conformation. PfMyoF contains two unique tail domains: a Rab-like and a WD40 domain, which together with its interactome and subcellular localisation support a role in membrane trafficking.

Although the PfMyoF amino acid sequence is highly divergent from human myosins, it retains all canonical motor domain features, including actin and ATP binding motifs, a conserved TEDS site, and an SH3-like N-terminal domain. Similar to PfMyoE, PfMyoJ and PfMyoK, PfMyoF contains a number of *Plasmodium*-specific insertions, particularly in the motor and converter domains. These large insertions most likely explain the reduced protein solubility when expressed in insect cells and their removal may improve expression yield for *in vitro* studies, as demonstrated for *P. falciparum* Kinesin-5^35^.

Within the neck domain we identify six IQ motifs that specifically bind PfCaM, the canonical calmodulin homologue in *P. falciparum*. These interactions stabilise the lever arm and support motor function. Processive motility within RPE cells suggests PfMyoF can also utilise human calmodulin, which shares 89% homology with PfCaM.

Using AlphaFold predictions, sequence alignments, and a live-cell filopodia assay, we establish PfMyoF as a plus-end-directed motor. Despite a large insertion within the converter domain, PfMyoF lacks Insert-2, a determinant of minus-end directionality, and aligns structurally with other plus-ended myosins. In contrast, PfMyoJ and PfMyoK retain Insert-2-like sequences consistent with minus-end motility, representing an important functional diversification in *Apicomplexan* parasites, which is absent from many other unicellular eukaryotes that do not possess minus-ended myosins, such as *Entamoeba* and *Saccharomyces*.

Directionality has important implications for PfMyoF function within parasites. Its plus-end directed, processive motility, together with its essential role in nutrient acquisition, suggests that PfMyoF may mediate the transport of vesicles through the parasite cytoplasm, likely towards the plasma membrane or organelles such as the Golgi complex ^36^. However, the organisation and orientation of actin filaments in *Plasmodium* remain poorly defined, although they are known to be highly dynamic^37^. A more detailed understanding of actin filament polarity within the parasite is therefore necessary to clarify the functional significance of PfMyoF directionality, particularly in the context of the apparent evolution of two minus-end-directed myosins, PfMyoJ and PfMyoK, in *Apicomplexa*.

Our analysis also explored dimerisation and autoinhibition of PfMyoF. Although AlphaFold-3-based modelling strongly predicted dimer formation via a coiled-coil domain, which was confirmed experimentally by enhanced tip accumulation in our cell-based assay, PfMyoF dimer formation has not yet been validated biochemically. Our results further suggest that PfMyoF can adopt a folded autoinhibited conformation through interactions between the N-terminal SH3-like domain and the WD40 domain. Indeed, full-length PfMyoF fails to localise to filopodia tips, consistent with an autoinhibited conformation, while deletion of either the SH3-like or the WD40 domain restores filopodia localisation and thus motility.

PfMyoF contains two unique, previously uncharacterised tail domains not found in HsMYO5A and other *Plasmodium* myosins. The Rab-like domain is a novel structural feature not previously seen in other myosin classes. It shares high sequence similarity with PfRab6 but lacks conserved Switch I/II motifs, suggesting a non-catalytic, pseudo-GTPase domain that may function in vesicle docking or Rab-effector recruitment. Whether it retains any GTP-binding capacity, and how this affects its function, remains an open question for future research.

The second PfMyoF tail domain, the seven-bladed WD40 β-propeller, strongly localises to the centrosome in human cells, consistent with centrosomal targeting of TgMyoF in *T. gondii* ^10^. Whether PfMyoF associates with and positions the centriolar plaque, the *P. falciparum* equivalent of centrosomes, remains to be determined ^38^. WD40 domains have also evolved in multiple *Apicomplexan* myosins, including TgMyoF, TgMyoH, TgMyoJ and TgMyoL, but are absent from non-*Apicomplexan* myosins, suggesting these domains have evolved independently to support parasite-specific functions.

Co-immunoprecipitation using a validated PfMyoF antibody identified several interactors involved in membrane trafficking, including PfRab18, dynamin-like proteins and actin regulators. These results further support a conserved role for PfMyoF in vesicle-associated processes, consistent with findings from *T. gondii,* where TgMyoF functions in organelle positioning and trafficking^31^. PfRab18, the most enriched interactor, is associated with vesicles destined for the parasitophorous vacuole membrane and exhibits perinuclear clustering with additional diffuse foci, a localisation pattern that is remarkably similar to endogenous PfMyoF^39^. Other interesting interactors include the myosin specific chaperone UNC-45, required for expression of soluble PfMyoA and PfMyoB, as well as centrin, a protein that could link the WD40 domain to the centrosome^40,41^. Whether CDPK7 phosphorylates the Rab-like domain of PfMyoF or is transported as cargo remains to be determined.

In summary, our findings establish that PfMyoF possesses a specialised protein architecture adapted for processive, long-range motility, supporting a central role in intracellular cargo transport in *P. falciparum*. PfMyoF appears to be the sole myosin dedicated to vesicular trafficking in this parasite, consistent with its proposed essential function. PfMyoF shares key features with HsMYO5A, including six IQ motifs bound to CaM that comprise a long lever arm capable of a large step size, plus-end directionality, likely dimerisation and SH3-like domain-mediated autoinhibition. This study characterises the key structural and functional features of PfMyoF, laying the groundwork for future investigations into cargo recognition, regulation by autoinhibition, and the roles of its unique Rab-like and WD40 tail domains. Our work provides a critical framework for future studies aimed at dissecting PfMyoF’s function and its potential as a therapeutic antimalarial drug target.

## Acknowledgements

We thank Dr Nicolas Bright for performing the immune electron microscopy. Mathew Gratian and Mark Bowen for help and advice with microscopy. Dr. Robin Antrobus and John Suberu for help and advice with proteomics. Sue Arden for molecular biology advice and thoughtful discussion, Theresa Feltwell for *Plasmodium* advice and training and Eva Pennink for comments on the manuscript. This work was supported in the Buss laboratory by a program grant from the Medical Research Council (MR/S007776/1), A.H. through an MRC University of Cambridge Doctoral Training Programme studentship (<u>MR/</u>N013433<u>/1</u>/<u>2430696</u>) and P.I. through a WACCBIP-World Bank ACE PhD fellowship (WACCBIP+NCDs: Awandare), CIMR microscopy is supported by equipment grants from the Wellcome Trust (108415) and the Medical Research Council (MR/Y002172/1). Work in the Rayner laboratory is supported by the Wellcome Trust (220266/Z/20/Z).

## Author contributions

AJH: conceptualisation, data curation, formal analysis, investigation, methodology, software, validation, visualization, writing – original draft and review & editing; PI: investigation, writing – review & editing; CB: investigation, methodology, writing – review & editing; AK: investigation, supervision, writing – review & editing; JCR: conceptualisation, resources, supervision, writing – review & editing; FB: conceptualisation, funding acquisition, project administration, resources, supervision, writing – review & editing.

## Methods

### Bioinformatics

#### Sequence Alignment

Sequences for PfMyoF (PF3D7_1329100), PfMyoJ (PF3D7_1229800) and PfMyoK (PF3D7_1140500) and other *P. falciparum* proteins were curated from PlasmoDB (https://plasmodb.org/plasmo/app). Multiple Sequence Alignments were performed using MUSCLE software (EMBL) and Pairwise Alignments performed using EMBOSS Stretcher (EMBL).

#### AlphaFold-3 Structural Prediction

AlphaFold Prediction and Structural Homology Protein structures were predicted using AlphaFold-3 software. Structures are coloured by the strength of prediction - with dark blue areas equalling high confidence and yellow/orange areas equalling low confidence. The Predicted Aligned Error (PAE) matrices shown can also be used to assess AlphaFold’s confidence in its prediction. The PAE plot represents a residue-by-residue error estimate matrix where both axes correspond to residue positions in the protein sequence. Darker green region in the matrix indicates high confidence interaction and lighter/white regions indicate low confidence. Squares of green indicate the presence of a protein domain. In addition, PAE plots are useful for examining predicted protein-protein interactions. To identify structural homologues, protein structures were saved as PDB files and uploaded to the DALI server for testing against both the PDB server and AlphaFold 2 databases for *H. sapiens* and *P. falciparum* and sorted by Z-score (a measure of similarity and significance).

For AlphaFold prediction of neck-light chain interactions, ten calmodulin and calmodulin-like proteins were identified using the PlasmoDB database: PfCaM (PF3D7_1434200); ELC (PF3D7_1017500); Putative CaM (PF3D7_1418300); Putative CaM (PF3D7_1030800); PF3D7_0714400 (PF3D7_0714400); PLC-3 (PF3D7_0414200); EF-hand calcium-binding domain-containing protein (PF3D7_1444200); EF hand domain-containing protein (PF3D7_0728500); calcium-binding protein (PF3D7_0605400); PLC-5 (PF3D7_0627200).

When predicted using AlphaFold, pTM scores ranged from 0.46 to 0.81, with only two proteins (PF3D7_0728500 and PF3D7_0714400) below the required >0.5 pTM score suggestive of a true-to-life model. Despite this, all putative light chains produced high confidence EF-hand domains responsible for IQ binding. PfCaM achieved a pTM score of 0.58.

### Cell Culture

#### RPE Cells

Human Cells Retinal Pigment Epithelial (RPE) cells were maintained in Dulbecco’s Modified Eagle’s medium (D6546, Sigma Aldrich) - HAM’s F-12 Nutrient Mix (N4888, Sigma Aldrich) supplemented with 2 mM L-glutamine, 100 U/mL penicillin 100 µg/mL streptomycin and 10% fetal bovine serum at 37°C with 5 % CO2. Cells were washed with phosphate-buffered saline (PBS, Sigma), incubated with 0.05 % trypsin-EDTA for a few minutes (Sigma) and transferred to a 15mL tube using in DMEM Buffer. Cells were passaged by dilution into fresh DMEM media in a new flask.

#### Transient Transfection

RPE cells were seeded onto glass coverslips in six well plates (typically 80,000 cells per coverslip), so that cells reached near confluency the following day. After settling and adhering to the coverslip, cells were transfected using FuGene® 6 transfection reagent as per manufacturer’s protocol. DNA (2 µg) and Fugene (6 µl) were made up to a total of 100 µl in Opti-MEM, mixed briefly, and incubated for 15 minutes at room temperature. The DNA-FuGene mix was then added dropwise to the cells.

#### *P. falciparum* Parasite Culturing

*Plasmodium falciparum* 3D7 cell lines were cultured in washed erythrocytes in modified RPMI media (*P. falciparum* 3D7 RPMI 1640 medium (R0883, Sigma-Aldrich) supplemented with 0.5% Albumax II, 0.2 mM hypoxanthine, 25 mM HEPES, sodium bicarbonate, and 12.5 µg/ml gentamycin). Cells were grown in a 37°C incubator 5 % CO_2_. During routine culturing, parasitaemia maintained below 5%. Parasitaemia was assessed every two days by Giemsa-smear. Parasites were synchronised by sorbitol lysis. Erythrocytes were pelleted by centrifugation and resuspended in 5% sorbitol. The cell suspension was vortexed for 15 seconds, incubated in a 37°C water bath for 5 minutes and vortexed again for 30 seconds to lyse erythrocytes infected with late-stage parasites. The remaining erythrocytes, those infected with ring stage parasites, were pelleted by centrifugation and resuspended in fresh media and restored to 4% haematocrit with washed blood.

### Protein Expression and Purification

#### *E. coli* Protein Expression

Proteins expressed in bacteria were cloned into a bacterial expression vector containing a tag for subsequent purification (either a 6-His tag in pRSETA or a GST tag pGex-4T). Expression vectors were transformed into BL21 cells, a single colony was picked and grown overnight in 5 mL 2xTY plus ampicillin at 37°C. Frozen stocks of the transformed bacteria culture were made by addition of 20% glycerol (1:1 ratio) and stored at -80°C. To express protein, a smear of frozen stock was inoculated in 50 mL 2xTY media plus ampicillin and grown overnight at 37°C at 200 RPM. The following day, the 50 mL culture was added to 1 L 2xTY plus ampicillin and grown at 37°C for 2-4 hours until the OD_600_ reached 0.6-0.8. OD_600_ values were measured on a Jenway 7305 UV/Visible Spectrophotometer. Once the required OD_600_ value was reached, the temperature was lowered to 22°C and culture cooled for 30 minutes. Cultures were then induced using 0.5 mM IPTG and grown overnight at 22°C at 200 RPM.

#### Sf9 Protein Expression

ExpiSf9 cells were maintained at 27°C, 125 rpm in *Spodoptera frugiperda* (Sf9) ExpiSf™ CD Medium (A3767801, Gibco™) at a density between1x10^6^ and 1x10^7^ cells/ml. Proteins were expressed from direct transfection of bacmid using ExpiFectamine or through viral infection. Coding sequences of the genes of interest were then optimised for expression in Sf9 cells and ordered as a g-block and cloned into pFastBac1 vectors. Constructs were cloned into bacmids, large plasmids containing genes for baculovirus expression, using BAC10 cells. Cells were transfected using ExpiFectamine SF transfection reagent (A38915, Gibco) according to the ExpiSf™ Expression System user guide (MAN0017532). Transfected cells were grown for 48-96 hours. Cells were pelleted at 310 x g for 5 minutes. The pelleted cells were stored or used immediately for protein purification.

#### Protein Purification

Cells were pelleted and supernatant discarded. The pellet was resuspended in lysis buffer (20 mM Tris pH 8.0, 500 mM NaCl, 10 mM MgCl_2_, 1 mM EGTA, 1 mM TCEP, protease inhibitors (Roche cOmplete Mini, EDTA-free), 0.1 % Triton X-100, 5 mM ATP), sonicated and centrifuged at 40,000 RPM at 4°C for 35 minutes in a Type 45 Ti rotor within an Optima XPN 90K ultracentrifuge. The resultant supernatant was carefully removed and filtered through a 0.22 µm filter. Glutathione Sepharose™ 4B (Cytiva) beads were washed three times in wash buffer, added to the cleared lysate and rocked end-over-end for 4 hours at 4°C. In the case of GFP-tagged proteins, a GST-GFP nanobody was also added to the cleared lysate. Beads were then pelleted by centrifugation at 2600 RPM for 5 minutes at 4°C and washed three times in wash buffer (20 mM Tris pH 8.0, 300 mM NaCl, 5 mM MgCl2, 1 mM EGTA) to remove nonspecific interactions. GST-tagged protein was eluted from using elution buffer (20 mM Tris pH 8.0, 300 mM NaCl, 5 mM MgCl2, 1 mM EGTA, 50 mM Reduced L-Glutathione) by incubating end-over-end for 20 minutes to 2 hours. The GST-GFP nanobody was removed from GST-tagged protein using PreScission Protease and the released PfMyoF isolated by incubation with Glutathione Sepharose™ 4B beads. Protein concentration was measured using the Precision Red Advanced Protein Assay (ADV02, Cytoskeleton) and proteins of interest identified by SDS-PAGE analysis.

#### SDS-PAGE

Denatured protein samples were resolved using Sodium Dodecyl Sulfate - PolyAcrylamide gel Electrophoresis (SDS-PAGE). Proteins were denatured by boiling for 5-10 minutes at 95°C in 5X sample buffer and loaded onto NuPAGE 4-12% Bis-Tris protein gels (NP0332) in NuPAGE 1X MOPS running buffer (NP0001) alongside Precision Plus Protein™ Standards ladder (Bio-Rad). Gels were run at a constant voltage of 200 V for 45 minutes at room temperature by an electrophoresis PowerPac™ (Invitrogen). Separated protein samples were visualised using InstantBlue™ Coomassie stain or processed for Western Blotting.

#### Western Blotting

Specific proteins of interest were identified by Western Blot. Proteins first separated by SDS-PAGE were transferred onto a nitrocellulose membrane or, for quantitative measures, a methanol-activated polyvinylidene difluoride (PVDF) membrane (Immobilon-P Milipore). Transfer took place in ice-cold 1X transfer buffer at 350 mA using an electrophoresis PowerPac™ (Invitrogen). Post-transfer, the membranes were blocked in 5% milk solution for 1 hour to prevent non-specific binding. The membranes were then incubated with primary antibodies diluted in PBST overnight before washing in PBST for 3x10 minutes. After incubating with secondary antibodies for 1.5 hours at room temperature and washing in PBST for 3x10 minutes, membranes were incubated with enhanced chemiluminescent (ECL) detection reagent (GE Healthcare) for 1 minute and exposed onto X-ray film (Fujifi).

#### Generation of Polyclonal Antibodies

GST-tagged PfMyoF WD40 fragments expressed and purified from *E. coli*, underwent size exclusion chromatography and were immunised by Eurogentec using their ‘classic’ antibody production programme alongside Freund’s complete adjuvant. The immunisation programme lasted 87 days in total.

#### Co-Immunoprecipitation and Protein Pulldown

100 ml of *P. falciparum* parasite cell cultures were grown to 5% parasitaemia and spun down for 5 minutes. Red blood cells were lysed by resuspension in 0.1% saponin made up in ice-cold PBS and complete protease inhibitors. After 5 minutes, remaining parasites were pelleted by spinning for 3 minutes at 3000 x g at 4°C. The pellet was resuspended in ice cold PBS + protease inhibitor and underwent repeated washes until all traces of heme/blood was removed. The parasite pellet was then resuspended in the lysis buffer (50 mM Tris pH 7.4, 150 mM NaCl, 5 mM MgCl2, 1% NP-40, 5 mM ATP, complete protease inhibitor (Roche cOmplete Mini, EDTA-free)) and incubated for 30 minutes on ice with occasional gentle pipetting. The cell lysate was passed through a 21G needle 3 times to shear DNA before centrifugation at 14,000 RCF to remove cellular debris. The resulting supernatant was incubated with Protein A end-over-end at 4°C for 45 minutes to remove protein non-specifically bound to the Protein A beads. Protein A beads were then added alongside 5 µg of antibody and incubated end-over-end for 90 minutes at 4°C. Beads were then pelleted and washed three times in the lysis buffer. All supernatant was removed with a U-100 insulin syringe, the remaining beads were then boiled in sample buffer and the purified protein sent for mass spectrometry.

### Microscopy

#### Light Microscopy

RPE Cells were fixed in 4% formaldehyde for 20 minutes, permeabilised in 0.2% Triton-100 for 2 minutes, blocked in 4% BSA/PBS for 60 minutes, and labelled with primary and secondary antibodies alongside Phalloidin and Hoechst. Coverslips were placed cell-side down onto ProLong Gold Antifade Reagent. Images were acquired using Zeiss AxioImager Z2 and LSM880 Airyscan confocal microscopes.

#### Filopodia Direction Assay

RPE cells were transfected with equal quantities of mCherry-tagged MYO6+ (a chimeric, plus-end directed MYO6) and GFP-tagged myosin constructs under investigation for directionality. After 20 hours, cells were fixed in 4% PFA and stained with anti-GFP, anti-RFP, Hoechst and Phalloidin-647. Coverslips were blinded prior to imaging using a Zeiss AxioImager Z2. For each GFP-tagged construct, 300 MYO6+ positive filopodia were analysed (100 filopodia per coverslip, across three coverslips). GFP signal was scored by eye as localising to the tip, shaft or being absent of filopodia based on reference to the archetypes established for MYO5A, MYO1E and MYO6 (Fig S4). Localisation percentages are presented in bar charts with standard error of the mean (SEM) shown.

#### Ultrastructural Expansion Microscopy (U-ExM)

Ultrastructural Expansion Micrsoscopy (U-ExM) and Iterative U-ExM (iU-ExM) typically results in a 5X and 20X enlargement of a microscopy sample respectively. All U-ExM and iU-ExM experiments were performed as detailed in [24, 29], respectively. In short, for U-ExM, washed *P. falciparum* infected erythrocytes were fixed in 4% formaldehyde for 30 minutes at room temperature. Fixed parasites were then allowed to settle onto poly-D-lysine coated 12 mm coverslips for 1 hour. Coverslips were incubated in 1 ml anchoring solution (1.4% Formaldehyde, 2% acrylamide, and Sodium acrylate 38% w/v in 1 X PBS) overnight at 37°C in 12 well dishes sealed with parafilm. The following day, a humid gelation chamber was assembled and placed in a -20 freezer 20 minutes prior to gelation. Monomer solution (19% w/v Sodium acrylate, 10% w/v Acrylamide, 0.1% w/v Bis (N,N’-methylenbisacrylamide)), 10% TEMED and 10% APS solutions were thawed in an ice-cold metal Eppendorf holder. Monomer solution (90 µl) was activated by addition of TEMED (5 µl) and APS (5 µl), quickly vortexed, and 35 µl pipetted onto the parafilm layer of the gelation chamber. The anchored sample was quickly blotted to remove excess anchoring solution and placed cell-face down onto the active monomer solution. Gelation tanks were left on ice for 5 minutes to allow complete absorption of solution by the cells and then placed at 37°C for 1 hour during which the gel is polymerised. For U-ExM, gels were boiled in 1.5 ml denaturation buffer (200 mM SDS, 200 mM NaCl, 50 mM Tris in pure water, pH 9.0) at 95°C for 30 minutes. After denaturation, gels were hydrated in ultrapure water and expanded overnight at 4°C. Water was completely replaced three times within the first hour of hydration to remove residual salt from the denaturation buffer. To immunolabel, 2x2 cm2 gels fragments were cut, placed in a 6-well dish and dehydrated by three 1X PBS washes and blocked in 3% BSA for 1 hour at 37°C. Primary antibodies were diluted into 3% BSA and incubated overnight in 37°C shaking rocker at 90 RPM. Gels were washed three times for 10 minutes in 1X PBST followed by addition of secondary antibodies and Hoechst diluted in 3% BSA. Gels were incubated with secondary antibody for a minimum of 5 hours shaking at 37°C. Gels were once again washed three times in PBST. For U-ExM, gels were hydrated overnight and ready for imaging.

Gels were imaged using Airyscan confocal microscopy on an LSM880 microscope. A small section of gel was cut and blotted extensively with Kimwipes. The gel was then gently pressed onto a poly-D-lysine coated coverslip using a paint brush. Once the correct plane of focus was found, pure water was added to the imaging dish to prevent gel dehydration during imaging.

#### Antibodies

α-PfMyoF (1/300, Rabbit, generated in this study); α-GFP (1/200, Rabbit, A-11122, Invitrogen); α-GFP (1/100, Mouse, 9F9.F9, Abcam); α-RFP (1/1000, Rat, Clone No. 5F8, Proteintech); α-γTubulin (1/250, mouse, Clone No. GTU-88, Santa Cruz); α-Acetylated Tubulin (1/250, Rabbit, 11H10, Cell Signalling Technology); α-Centrin (1/250, Mouse, Clone No. 20H5, Sigma), α-PvERD2 (1/500, Rabbit, gifted from the Rayner Lab), α-PvBiP (1/250, Rat, gifted from the Rayner Lab), Alexa Fluor-conjugated secondary antibodies (1/1000, Donkey & Goat, Invitrogen); Phalloidin 647 (1/1000, A22287, Invitrogen); Hoechst (1/1000, 33342, Invitrogen); Atto 565 NHS Ester (1/300, 72464, Sigma).

#### Immunoelectron microscopy

Parasites were tightly synchronised to a 4-hour window by repeated rounds of sorbitol and percoll synchronisation. The egress inhibitor Compound 2 was used to arrest mature schizonts for 4 hours prior to Percoll purification and fixation with 4% paraformaldehyde / 0.075% glutaraldehyde in PBS for 30 minutes at 37 °C. The cell suspension was then pelleted (13 000 rpm for 5 min), the fixative was aspirated, and the cell pellet was re-suspended in warm 10% gelatin. The cells were then pelleted (13 000 rpm for 5 min) and the gelatin-enrobed pellet was set on ice, trimmed into 1mm3 blocks and infused with 2.3M sucrose for 24h at 4°C. The blocks were subsequently mounted on cryostubs and snap-frozen in liquid nitrogen. Frozen ultrathin sections (70 nm) were cut using a Diatome diamond knife in a Leica FC7 cryochamber attachment (Leica, Milton Keynes, UK), collected from the knife-edge with 50:50 2% methyl cellulose: 2.3 M sucrose and mounted on formvar-carbon coated EM grids.

Immunolabelling was performed using the protein A-gold technique at room temperature^42^. Sections were incubated with 50 mM NH_4_Cl in PBS for 10 minutes to quench unreacted aldehydes and then 1% BSA/5% FCS in PBS for 10 minutes. Sections were incubated for 30 minutes with 5 ml of PfERD2 antibody 1:50 in PBS containing 5% FCS and 0.1% BSA. The sections were washed with PBS/0.1% BSA (6 x 3 minutes) and incubated for 30 minutes with PBS/0.1% BSA containing protein A conjugated to 10 nm colloidal gold. The sections were washed with PBS/0.1% BSA (2 x 5 minutes), PBS (4 x 5 minutes) and the complex stabilised using 1% glutaraldehyde in PBS (5 minutes).

Sections were incubated with 50 mM NH_4_Cl in PBS for 10 minutes to quench unreacted aldehydes and then 1% BSA/5% FCS in PBS for 10 minutes and were then incubated for 30 minutes with 5 ml of PfMyoF antibody diluted 1:50 in PBS containing 5% FCS and 0.1% BSA. The sections were washed with PBS/0.1% BSA (6 x 3 minutes) and incubated for 30 minutes with PBS/0.1% BSA containing protein A conjugated to 15 nm colloidal gold. The sections were washed with PBS/0.1% BSA (2 x 5 minutes), PBS (4 x 5 minutes). Finally, the sections were rinsed with distilled water (5 x 3 minutes) and contrasted by embedding in 1.8% methyl cellulose / 0.3% uranyl acetate. Sections were allowed to air dry prior to observation in an FEI Tecnai G2 BioTWIN transmission electron microscope (FEI, Eindhoven) at an operating voltage of 80kV. Images were recorded with an Eagle 4K CCD camera.

