## Supplementary Figure 1-5 for "Plasmodium falciparum Myosin F with a Rab-like tail domain localises to perinuclear membranes and associates with trafficking proteins"

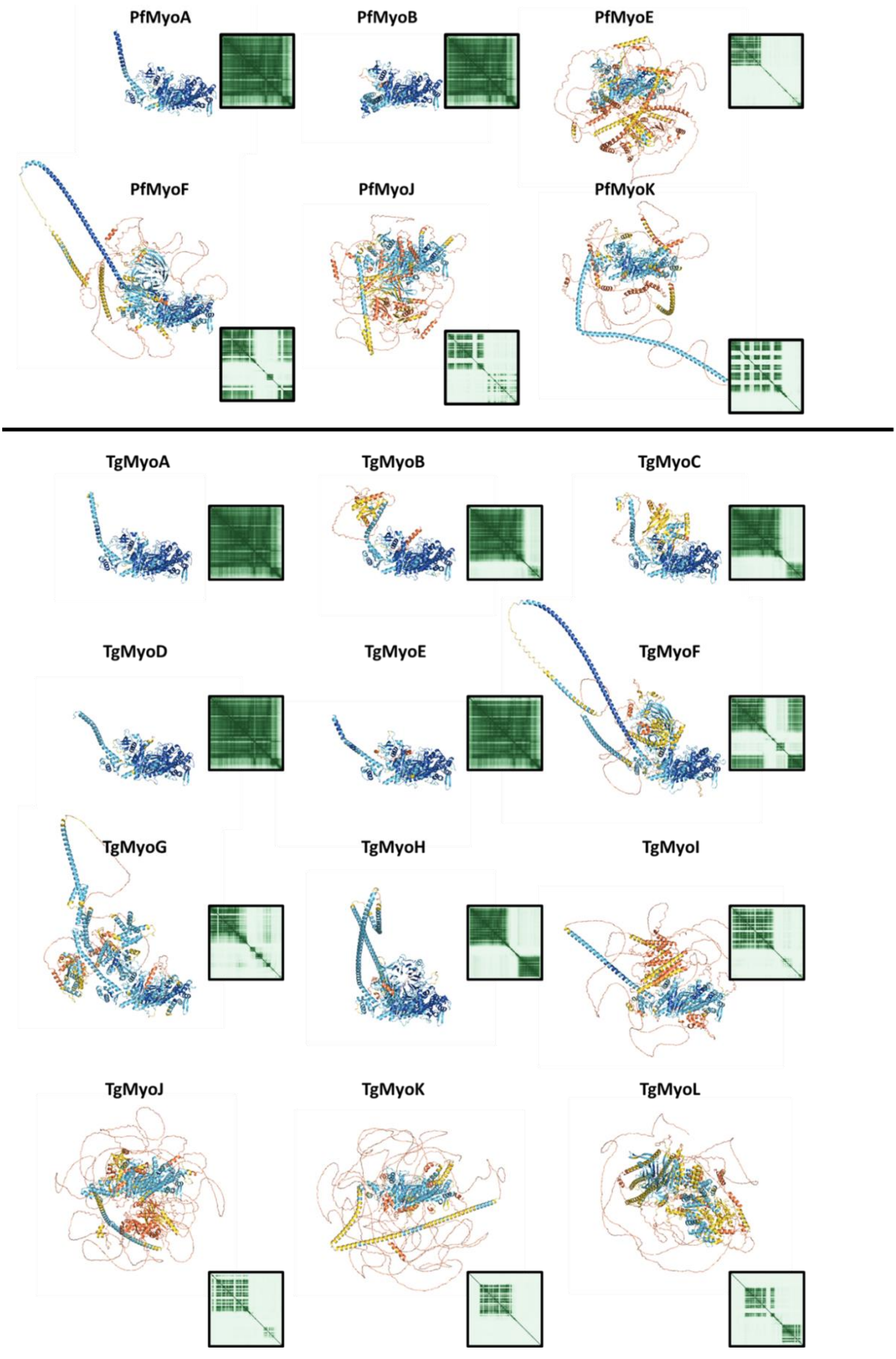


Supplementary Fig 1 – AlphaFold-3 predictions of the six *P. falciparum* myosins (top) and eleven *T. gondii* myosins (bottom). PAE graphs are shown to the right of each structure.

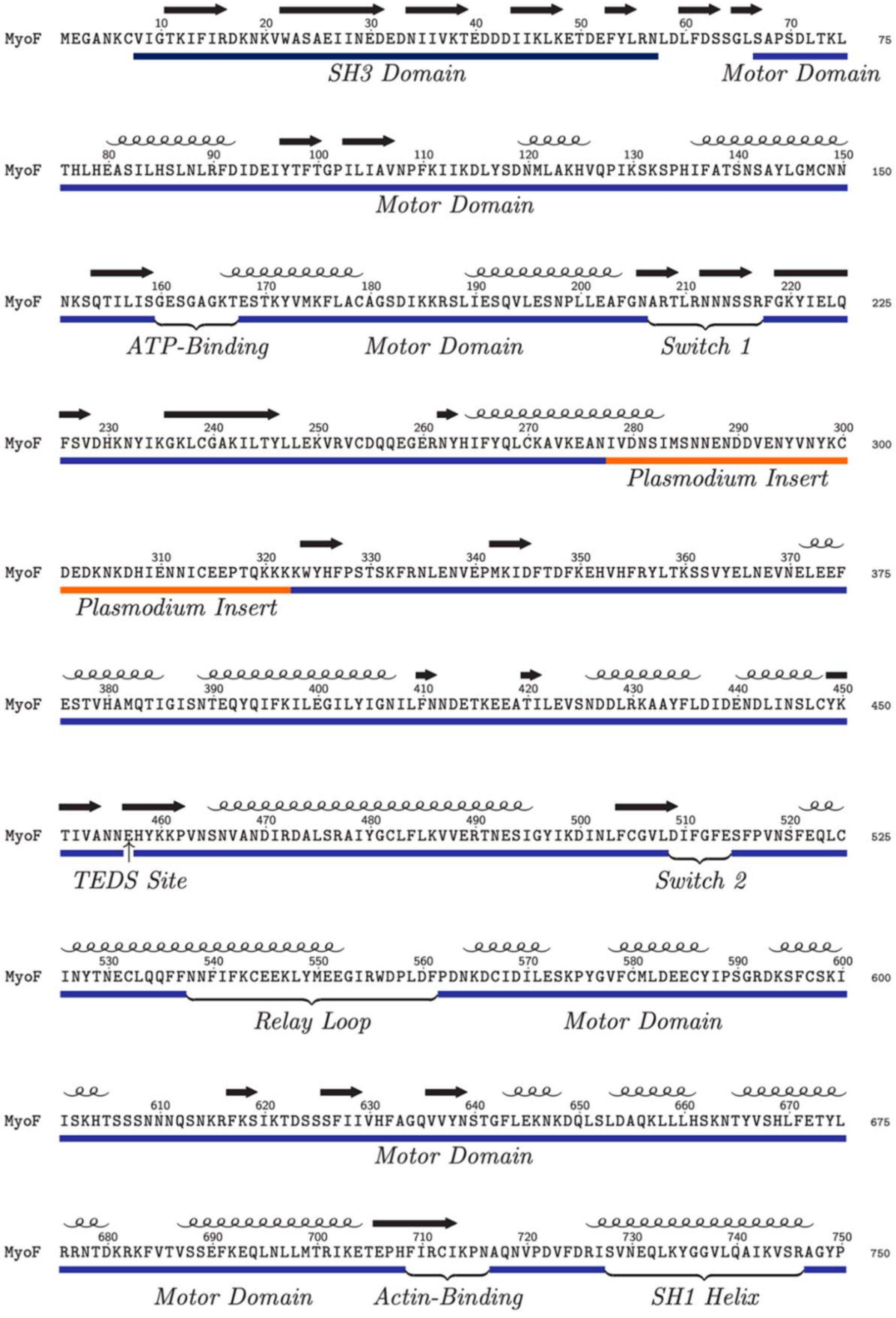


Supplementary Fig 2A – The Annotated PfMyoF Motor domain. Alpha helices are labelled as such and β sheets as black arrows. Orange sequences highlight large asparagine-rich Plasmodium specific insertions

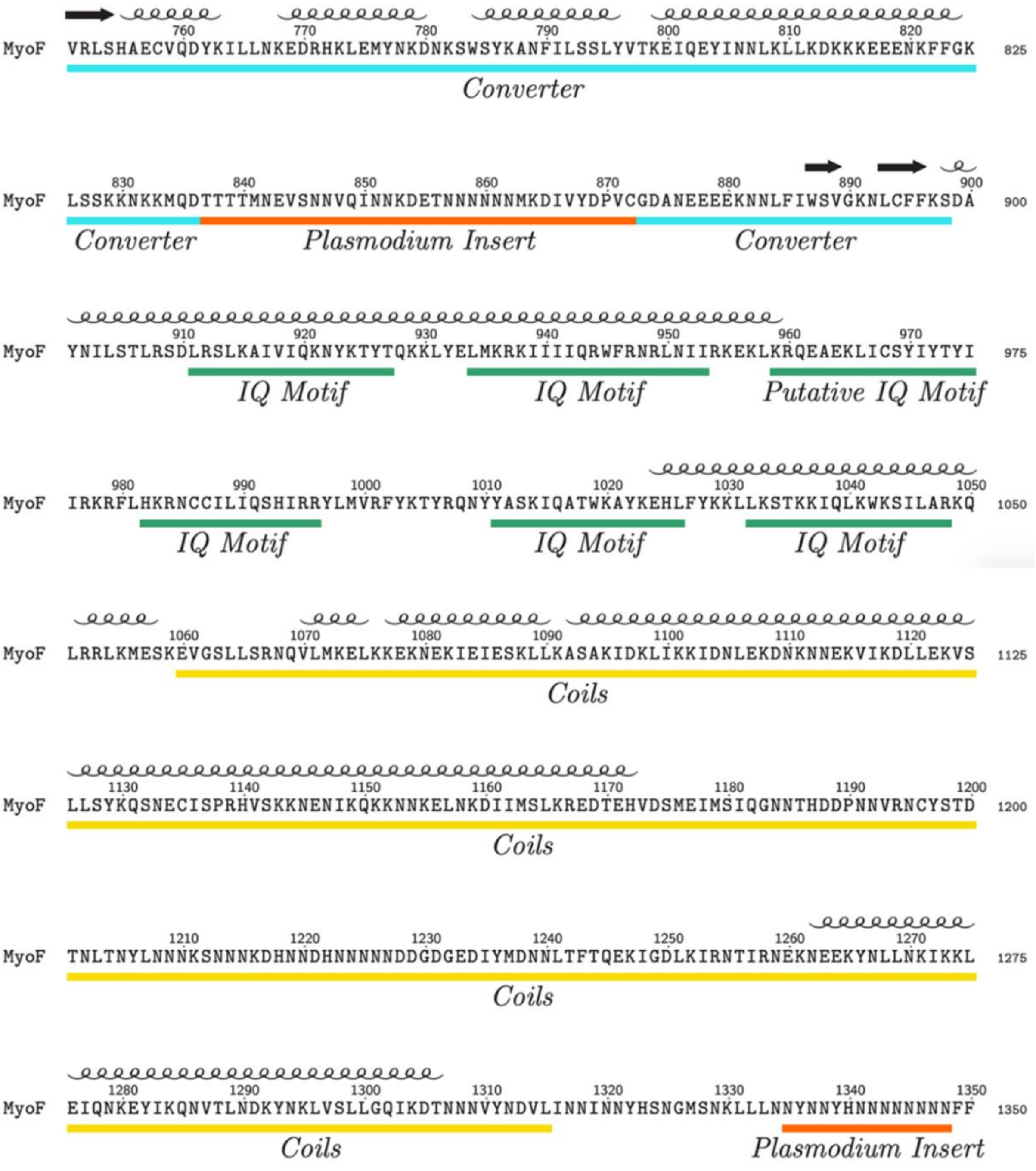


Supplementary Fig 2B – The Annotated PfMyoF Converter, Neck and Coil domains. Alpha helices are labelled as such and β sheets as black arrows. Orange sequences highlight large asparagine-rich Plasmodium specific insertions

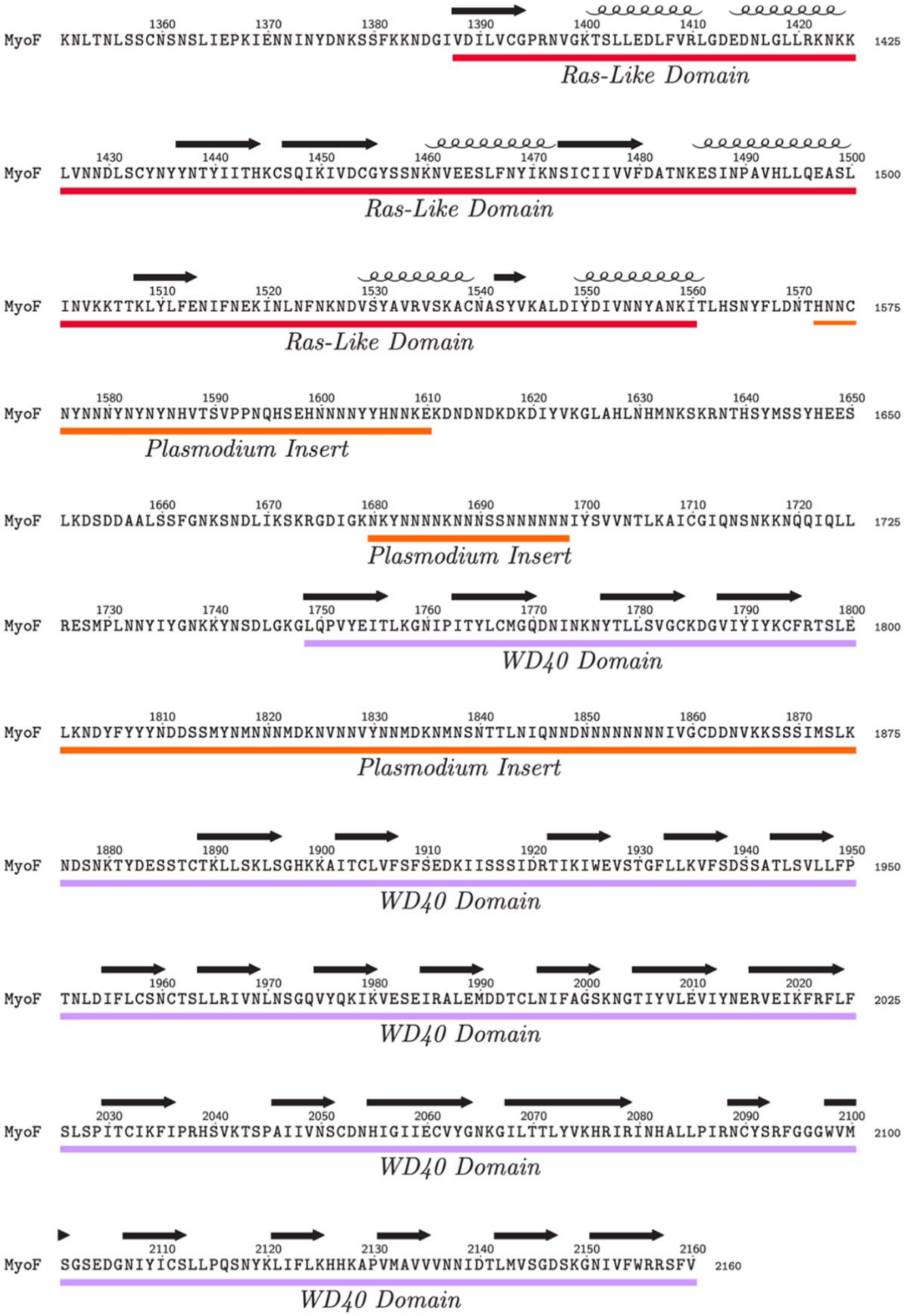


Supplementary Fig 2C – The Annotated PfMyoF Tail domain. α-helices are labelled as such and β-sheets as black arrows. Orange sequences highlight large asparagine-rich Plasmodium specific insertions

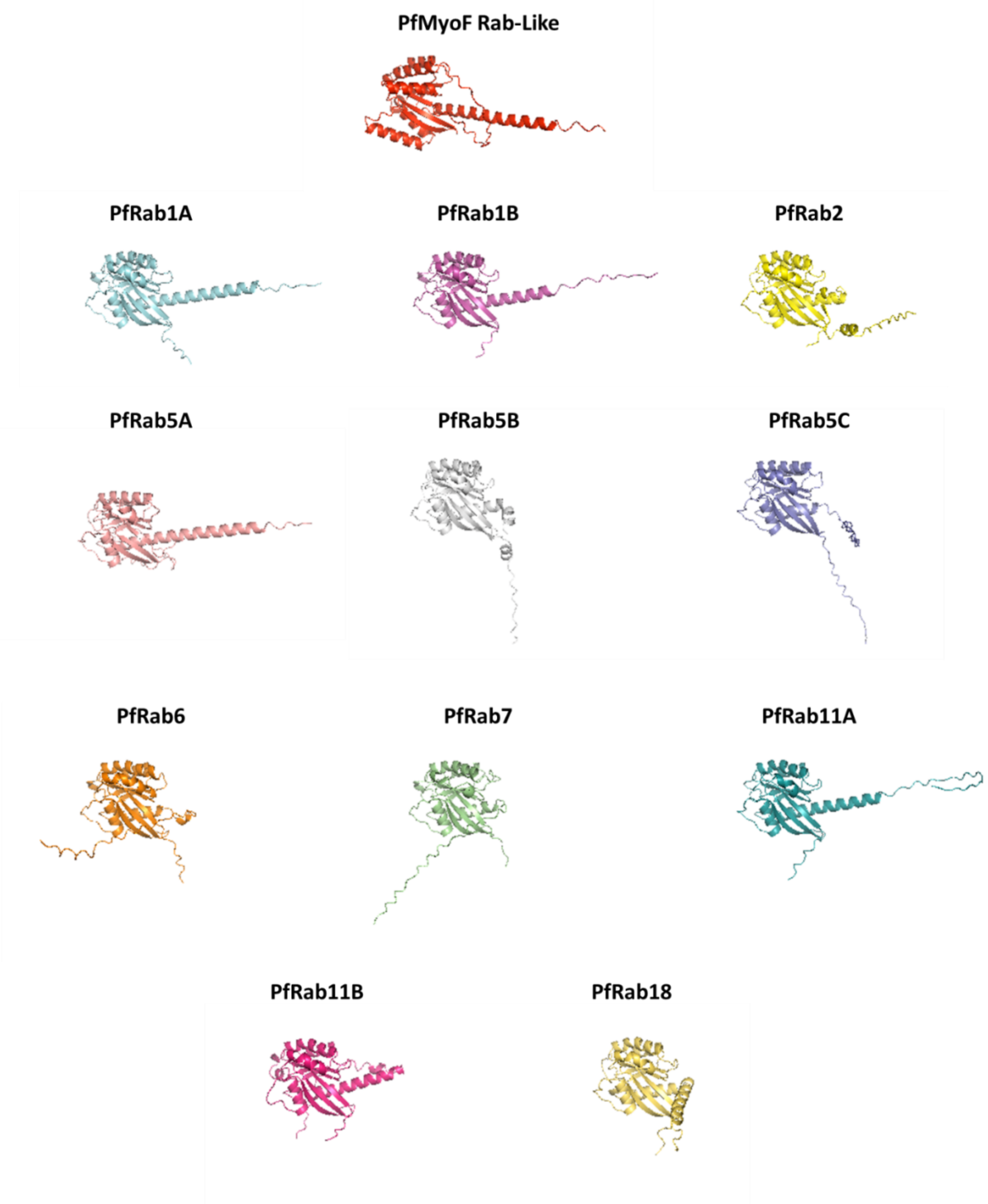


Supplementary Fig 3 – AlphaFold-3 predictions of the PfMyoF Rab-Like domain alongside the 11 Rab proteins of *P. falciparum*, revealing high structural similarity.

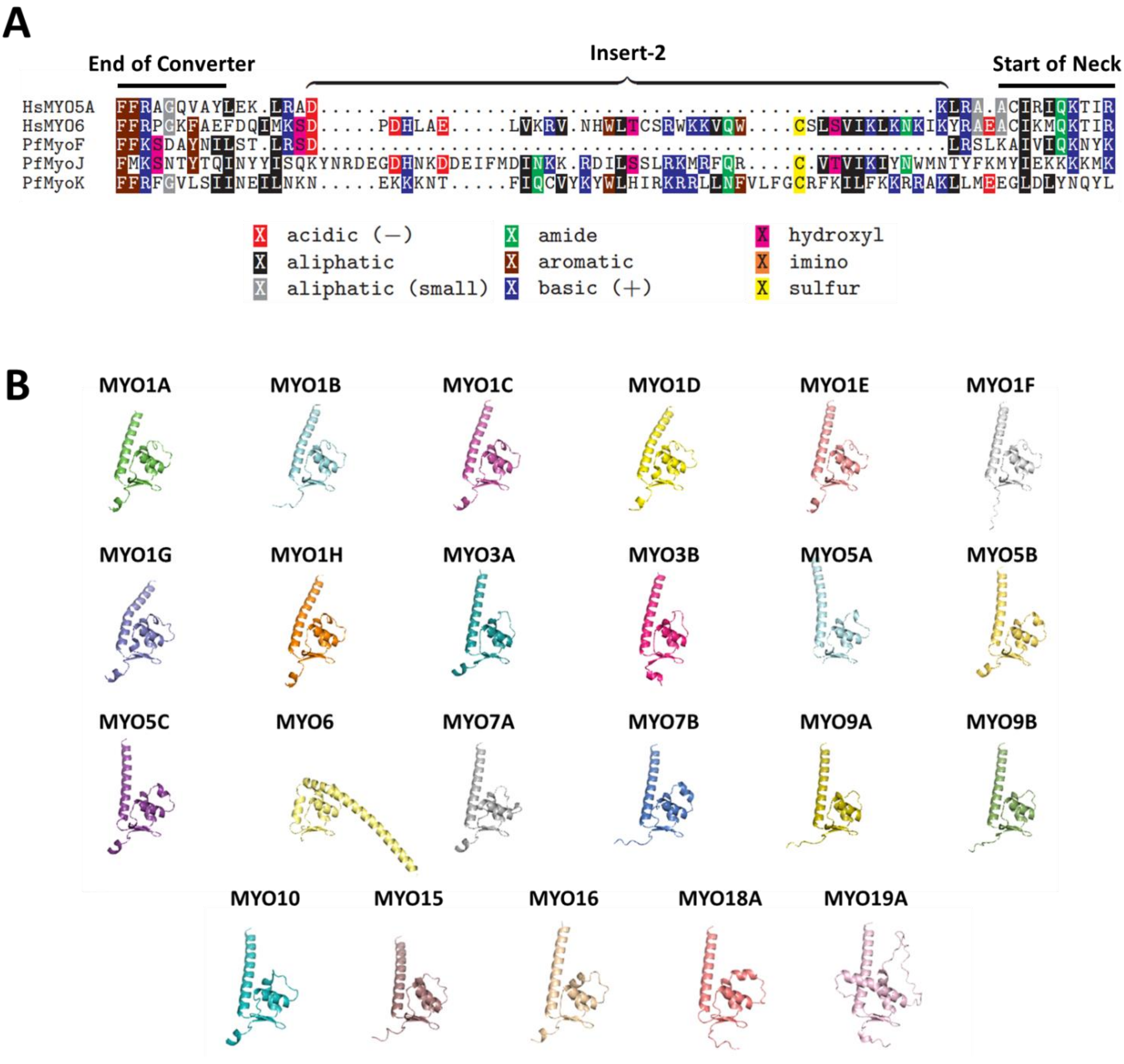


Supplementary Fig 4 – (A) Multiple Sequence Alignment of 'Insert-2' regions at the end of the converter domains of HsMYO5A, HsMYO6, PfMyoF, PfMyoJ and PfMyoK. Both HsMYO5A and PfMyoF lack Insert-2, which is present in PfMyoJ and PfMyoK. PfMyoJ contains a sequence strong with strong homology to the Insert-2 of HsMYO6. (B) AlphaFold-3 prediction and alignment of the converter-neck angles of all unconventional human myosin proteins, which identifies HsMYO6 as a minus-end directed motor based on lever arm position.

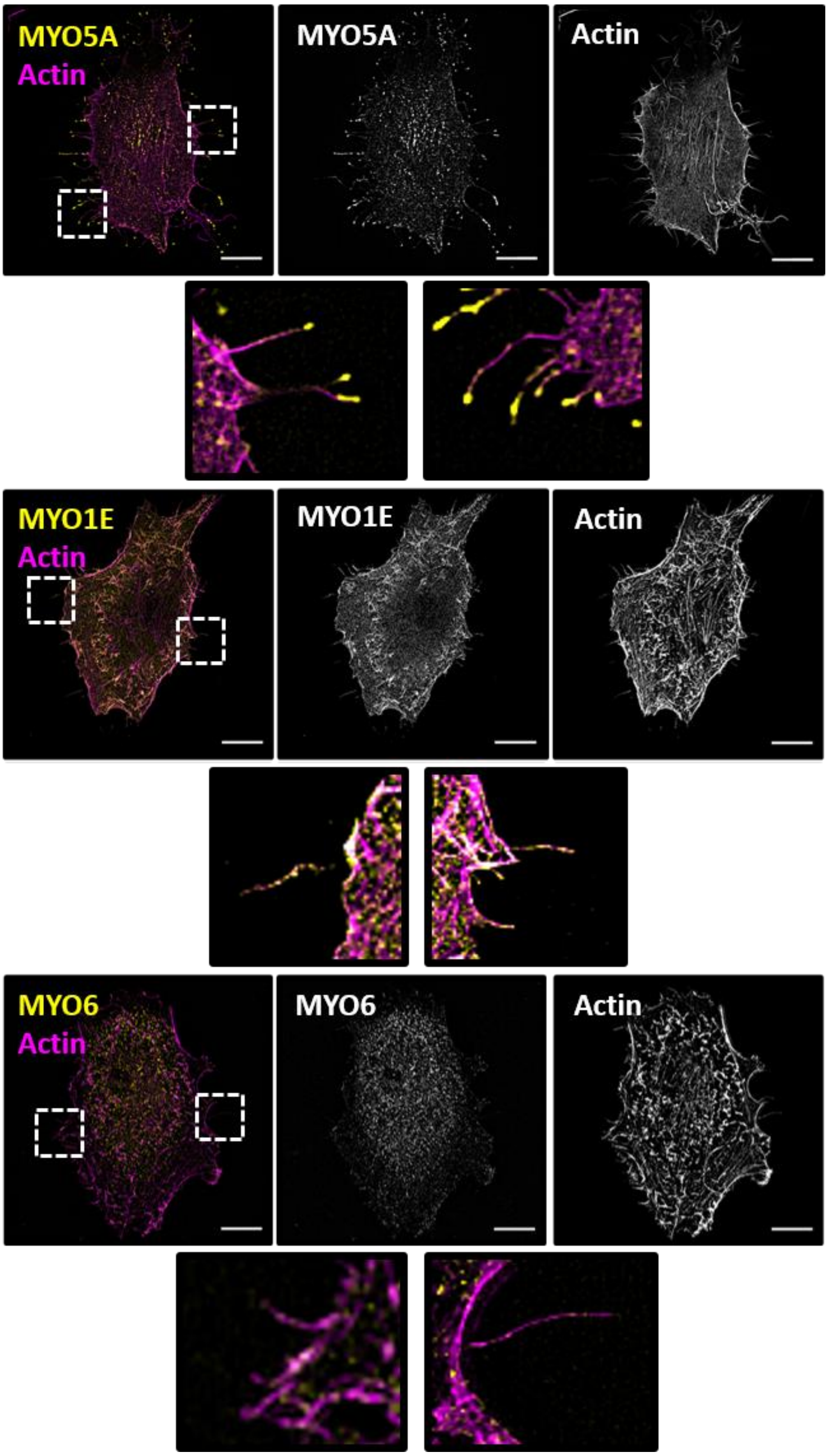


Supplementary Fig 5 – The Archetypes of Filopodia Localisation. Representative images of three types of myosin localisation in filopodia of RPE cells. Tip end localisation (MYO5A), shaft localisation (MYO1E) and absent (MYO6). Actin (magenta) and GFP-tagged myosin (yellow) are shown. Scale bar, 10 μm.
